# Biosynthesis of glycosylated 5-hydroxycytosine in the DNA of diverse viruses

**DOI:** 10.1101/2025.06.16.659550

**Authors:** Yan-Jiun Lee, Jianjun Li, Cecilia Sobieski, Sophie H. Young, Jacek Stupak, Evguenii Vinogradov, Hongyan Zhou, Wangxue Chen, Peter R. Weigele, Danielle L. Peters

## Abstract

Characterizing viral strategies for circumventing cellular defenses is critical to understanding the biology and therapeutic application of bacteriophages as antimicrobials. Many bacteriophages synthesize complex modifications (i.e., hypermodifications) to the nucleobases of their virion DNA in order to circumvent the endonuclease-based defenses of their hosts. To date, most hypermodified pyrimidines in viruses are synthesized via group transfer to a pre-existing hydroxyl moiety in DNA during the later stages of lytic development leading up to packaging of the viral chromosome into the capsid. Typically, these occur at hydroxymethylcytosine or hydroxymethyluracil. We find mono- and poly-arabinosylated cytidines completely replacing canonical cytidine in the native DNA of the Escherichia phage RB69 and *Acinetobacter baumannii* phage DLP3. In both cases, an arabinosyl moiety is connected directly to the pyrimidine through an ether linkage with the bridging oxygen originating from 5-hydroxycytosine (5hoC) in DNA. The 5hoC is synthesized as a mononucleotide by a virally encoded thymidylate synthase homolog. We find diverse virally encoded thymidylate synthase homologs, including the *Rhizobium* phage RL38J1, are capable of producing 5hoC suggesting this base modification is a starting point for glycodiverse cytosine derivatives. Characterizing these cytosine derivatives adds to our understanding of viral anti-defense mechanisms.

**GRAPHICAL ABSTRACT:** 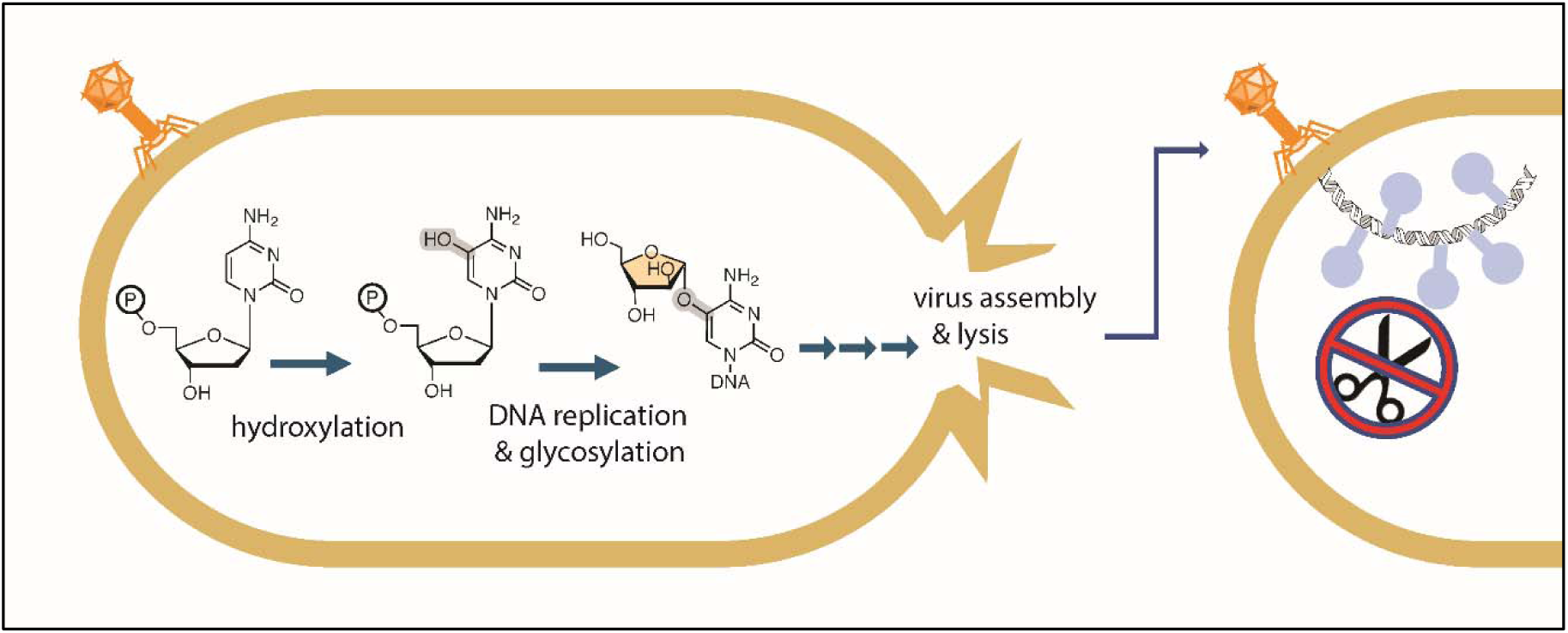

## INTRODUCTION

Viruses of bacteria (bacteriophages) have evolved a myriad of ways to evade the defense mechanisms of their cellular hosts, including small protein inhibitors of restriction endonucleases (1), anti-CRISPR proteins (2), specialized subcellular compartments for viral morphogenesis (3, 4), and complex non-canonical nucleobases known as DNA hypermodifications (5). Given the ongoing discovery of new anti-phage systems as well as novel virally encoded anti-defense systems, it is likely that for any given bacteria-encoded defense system, there exists in the virosphere a bacteriophage encoded mechanism evolved to overcome it.

Because of their ubiquity, host-specificity, and diversity, bacteriophages represent a vast and largely untapped reservoir of potential antimicrobial therapeutics. Understanding the diverse mechanisms by which they can overcome bacterial cells is imperative to their engineering and application to treating infections (6). Antimicrobial resistant (AMR) bacterial infections are a troubling global health crisis, with 1.14 million deaths attributed to AMR in 2021– a number forecasted to increase to 1.9 million by 2050 (7). Due to the increase in AMR infections, and significant lack of research into new antibiotics, alternatives to traditional antimicrobials are required to address this urgent silent pandemic (8). The top priority pathogens highlighted are Gram-negative carbapenem-resistant bacteria, such as Carbapenem-Resistant *Acinetobacter baumannii* (CRAb) and *Escherichia coli* (9). How these pathogens might resist infection by bacteriophages is an important consideration for designing a bacteriophage therapeutic against them.

Bacteria and archaea commonly employ endonuclease effectors as part of restriction-modification (RM) innate immunity systems and CRISPR-Cas adaptive immunity systems to defend against bacteriophages and other mobile genetic elements. In response, many bacteriophages have evolved alternative nucleobases in their DNA which are resistant to recognition and/or cleavage by endonuclease-based defenses. Collectively, the Caudoviricetes (i.e. “tailed” dsDNA viruses of bacteria and archaea) are revealing a growing repertoire of chemically distinct nucleobases functioning in place of each of the four canonical nucleotides. DNAs containing hypermodified pyrimidines have been shown to be resistant to the majority of commercially available Type II restriction endonucleases in vitro (10, 11) and the glucosylated cytidines of the T-even bacteriophages have been shown to inhibit CRISPR-Cas systems (12, 13). On the other hand, hypermodified DNA exposes bacteriophages to restriction by Type IV (modification dependent) restriction endonucleases (14–16) and defense glycosylases (17). Thus, understanding the genetics and biochemistry of pyrimidine hypermodification may contribute to a predictive understanding of the therapeutic potential of a given bacteriophage against pathogenic hosts of a certain genetic background.

All currently known hypermodified deoxypyrimidines bear moieties at C5 of the base heterocycle which are different to their canonical counterparts (18). Cytosine hypermodification in bacterial viruses typically involves the formation of a hydroxymethylated precursor in DNA followed by group transfer of a sugar from a nucleotide diphosphate carrier to the hydroxymethyl at C5, catalyzed by DNA glucosyltransferases (18). Recently, it has been demonstrated that 5-hydroxymethylcytosine (5hmC) can be synthesized in DNA through oxidation of 5-methylcytosine (5mC) by bacteriophage encoded homologs of the Ten-eleven-translocation (Tet), with the modified bases subsequently glycosylated by virus encoded transferases (19). However, most cultured viruses containing hypermodified deoxypyrimidines (e.g. contractile-tailed bacteriophages belonging to the *Straboviridae*, *Herelleviridae*, *Mesyanzhinovviridae*, and *Ackermanviridae*) produce non-canonical pyrimidine mononucleotides such as 5-hydroxymethyl-2’- deoxyuridine monophosphate (5hmdUMP) and 5-hydroxymethyl-2’-deoxycytidine monophosphate (5hmdCMP) using bacteriophage encoded homologs of thymidylate synthases (18). These non-canonical nucleotides are sequentially converted to their triphosphate forms through viral and host mononucleotide kinases and incorporated into DNA by viral replicative DNA polymerases. Non-canonical deoxypyrimidine biosynthesis is often accompanied by parallel pathways that diminish or eliminate the corresponding canonical deoxypyrimidine, leading to its complete replacement in the viral DNA (5). Prior to being packaged into a capsid, many bacteriophage DNAs are further modified with sugars, amino acids and other small molecules by transferases resulting in hypermodified DNA (5, 20).

The pentose arabinose was recently identified as a substituent of modified cytidines in the DNA of coliphage RB69 and the *Shewanella* phage Thanatos (21). However, the complete chemical structure of the arabinosylated cytidine was not reported. While investigating the hypermodified cytidines of coliphage RB69, as well as the CRAb infecting *Acinetobacter* phage DLP3, we determined the modification as arabinose conjugated directly to the cytosine heterocycle at position 5 via a bridging oxygen. Interestingly, we find the arabinosylated cytosine to be derived from a hydroxylated cytidine produced at the mononucleotide level by bacteriophage encoded thymidylate synthase homologs, an activity unexpected for an enzyme normally associated with transfer of a single carbon (EC 2.1.-.-, enzymes transferring a single carbon).

## MATERIAL AND METHODS

### *Acinetobacter baumannii* phage DLP3 propagation and DNA isolation

The DLP3 host is a capsule mutant of *A. baumannii* AB5075 (AB5075cm), generously provided by Dr. Sebastien Crepin, who acquired the strain from the Manoil lab. The host was grown aerobically overnight at 37 °C on Lennox Luria Bertani (LLB) agar plates or in LLB broth with shaking at 200 rpm.

Phage DLP3 was isolated from an Ottawa, Ontario sewage sample using previously described methods (22). To prepare phage lysate for DNA extraction, a subculture of AB5075cm was supplemented with 5 mM CaCl_2_ and MgCl_2_ and grown to an OD600 of ∼1.2. Next, DLP3 was added at a final concentration of 1x10^8^ PFU/mL and incubated at 37°C for 18 hrs with 200 rpm shaking. The culture/phage lysate was centrifuged at 12,000 × g for 10 min at 4 °C to pellet the bacterial host. The cleared phage lysate was filter-sterilized with a 0.22 µm PES filter (Millex Cat. No. SLGPR33RB) and then stored at 4 °C until further use.

A nuclease digestion buffer, DNase I (1 U/µl, ThermoFisher, Cat.# AM2222) and RNase A (100 mg/ml, Qiagen Cat.# 19101) were added to the DLP3 phage lysate and incubated at 37 °C for 30 min to degrade exogenous DNA and RNA. Next, ZnCl_2_ was added to the lysate at a final concentration of 40 mM, followed by incubation at 37°C for 5 min to precipitate the phage. The sample was centrifuged at 8,000 x g for 10 min, and the resulting pellet was resuspended in 1x PBS. Next, EDTA, SDS, and Proteinase K (Promega Cat.# MC5005) were added to the lysate at final concentrations of 0.05 M, 1 %, and 0.5 mg/mL, respectively. The sample was gently mixed for 3 min, and incubated at 55°C for 60 min. Next, the lysate was mixed three times with an equal volume of phenol-chloroform isoamyl alcohol (PCI, ThermoFisher Cat.# 15593031), then an equal volume of chloroform, and the aqueous phase recovered each time. The aqueous phase was mixed with 1/10 volume of 3 M sodium acetate pH 5.2 and 2.5 volumes of 99% ice-cold ethanol to precipitate the DNA. The DNA was then washed thrice with 70 % ethanol, air dried, and resuspended in nuclease-free water. The DNA was digested to nucleosides using the Nucleoside Digestion Mix (NEB, Cat.# M0649S) following the manufacturer’s supplied protocol.

### *Escherichia coli* phage RB69 propagation and DNA isolation

RB69 infects general laboratory *E. coli* K and B strains. For the presented work, a derivative of the *E. coli* K strain T7 Express (NEB, Cat# C2566) was used, and it was grown aerobically at 37 °C on LLB broth with agitation overnight.

Phage RB69 was obtained from Prof. Jim Karam of Tulane University. To propagate phage, a subculture of T7 Express with an OD600 of ∼0.1 was infected with RB69 plaques and incubated at 37°C for 10 hr with 250 rpm shaking. The culture was centrifuged at 12,000 × g for 10 min at 4 °C. DNase I (NEB, Cat# M0303) and RNase A (NEB, Cat.# T3010L) were added to the phage solution at 100 µg/mL and 10 µg/mL, respectively, then incubated at 37°C for 30 min with gentle swirling. Followed by adding solid NaCl and poly(ethylene glycol) (average MW. 8000) at a final concentration of 1 M and 10% (w/w), respectively, and incubated at 4°C for 4 hrs to overnight; the phages were then pelleted by centrifugation at 12000 x g off at 4°C for 10 min. SM buffer (50 mM Tris-HCl pH 7.5, 100 mM NaCl, 8 mM MgSO_4_, 0.01% Gelatin) was used to resuspend the phage pellet, and the phage solution was stored at 4°C in the dark until further use.

To extract the RB69 genomic DNA, EDTA, SDS, and proteinase K (NEB, Cat# P8107) were added to the lysate at final concentrations of 0.05 M, 1 %, and 0.5 mg/mL, respectively. The sample was gently mixed and was incubated at 55°C for 30 min or until the solution appears to be translucent. Next, a phenol-chloroform extraction was performed where the sample was mixed three times with an equal volume of phenol-chloroform isoamyl alcohol (VWR, Cat.# 97064-716), then an equal volume of chloroform, and the aqueous phase recovered each time. The aqueous phase was mixed with 1/10 volume of 3 M sodium acetate pH 5.2 and 2.5 volumes of 99% ice-cold ethanol to precipitate the DNA. The DNA was then washed thrice with 70 % ethanol, air dried, and resuspended in nuclease-free water.

### Restriction digestion of DLP3 and RB69 genomic DNA

All restriction enzymes used are from New England Biolabs, Inc., (NEB). Approximately 200 µg of purified DLP3 or RB69 genomic DNA was used in each restriction digestion reaction. 20 U of each enzyme was added into the DNA solution in 1X CutSmart buffer in a volume of 25 µL and the reaction was incubated at 37°C for 2 hrs then heat-inactivated at 65°C for 20 minutes. Digested DNAs were analyzed by agarose gel electrophoresis using 1.2% agarose gel in 1X TBE buffer with 1 µg/mL ethidium bromide. After electrophoresis, the gel was visualized under ultraviolet exposure at 260 nm and the gel image was documented by using a CCD camera. A virtual digestion for each restriction enzyme used was generated using NEBcutter v2, assuming the DNA contains only canonical bases. The experimental DNA cleavage results were compared to the virtual digestion pattern to assess the resistance of restriction by the base modification in the DNA.

### LC-MS methods for hydrolyzed phage virion DNA

Approximately 0.5 µg DLP3 or RB69 phage DNA was hydrolyzed and dephosphorylated using the Nucleoside Digestion Mix (NEB, Cat.# M0649) following the manufacturer’s protocol at 37°C for 14 hrs. The resulting nucleosides mixture was filtered through a hydrophilic PTFE 0.2 µm centrifugal filter, and the filtrate was subjected to reverse-phase HPLC-MS analysis.

LC-MS was performed on an Agilent LC-MSD XT system configured with a 1290 Infinity II LC and a G6135 single quadrupole mass detector. A Waters XSelect HSS T3 C18 column (2.1 × 100 mm, 2.5 µm particle size) was used for reverse-phase liquid chromatography, which was conducted in a linear gradient from 1 to 10% in 10 min of binary mobile phases composed of 10 mM ammonium acetate (pH 4.5) and methanol operated at a 0.6 mL/min flow rate. The chromatography was monitored at 260 nm UV absorbance. For the MS detection, the source electrospray ionization was operated in positive (+ESI) and negative (-ESI) modes. MS acquisition was performed with a capillary voltage of 2500 V at both modes, a 70 V fragment voltage, and a mass range of m/z from 100 to 1000. Agilent ChemStation software was used to process the primary LC-MS data. ChemStation-generated chromatograms were further annotated in Adobe Illustrator.

### LC-MS/MS of hydrolyzed phage DNA

The phage DNA was first digested to nucleosides as described in the LC-MS section. The afforded aqueous solution was passed through SepPak C18 column, washed with water, and then methanol where the methanol wash fraction contained a mixture of the nucleosides. A Zorbax C18 column was used to further purify the mixture.

Reverse phase LC-MS/MS analysis of nucleosides was performed on an Exploris Orbitrap 240 mass spectrometer coupled to an Ulitmate3000 capillary HPLC (Thermo Scientific, Toronto, Canada). Data-dependent acquisition (DDA) was carried out in the positive-ion mode. Liquid chromatography separation was performed using Waters Atlantis T3 columns (150 x 1 mm; 130 Å, 3 μm) with a 35 µL/min flow rate and the following gradient program: 0 - 1 min 0 % B, 1 - 12 min increasing to 60 % B, 12 – 17 min 60 % B, 17.1 min 90 % B held until 21 min, and equilibration from 21.1 – 35 min. Mobile phase A contained aqueous 2 mM ammonium bicarbonate, and mobile phase B was methanol. Mass spectrometer operated in the positive-ion mode and the electrospray voltage was kept at 2800 V to prevent reduction of DNA bases in the electrospray source.

### Absolute configurations analysis using GC/MS

The method for the determination of absolute configurations has been reported by others (23). Briefly, the samples (∼0.5 mg) (R)- or (RS)-2-octanol (0.2 mL) and acetyl chloride (0.02 mL) were added at room temperature, heated in closed vials (100 °C, 2 hrs), dried by air stream, acetylated (0.2 mL Ac_2_O, 0.2 mL pyridine, 100 °C, 30 min), dried, then analyzed by GC-MS on Thermo Trace 1310 instrument with ITQ1100 ion trap detector, capillary column HP-5 at 160-260 °C by 4 °C/min temperature gradient.

### Modified nucleoside sample preparation for NMR spectroscopy

A two-step workflow composed of solid-phase extraction followed by liquid chromatography was used to enrich and prepare the modified nucleoside samples from hydrolyzed DLP3 and RB69 phage DNA for NMR spectroscopy. Briefly, the hydrolyzed DNA isolated from phages was diluted in water before being applied to a SepPac C18 cartridge pre-conditioned by washing with methanol (10 mL) and water (10 mL). 70% methanol or 100% methanol (∼5 mL) were used to elute DLP3 or RB69 DNA samples from the column, respectively. The methanol fraction was confirmed to contain the desired compound with mass spectrometry. For DLP3 samples, the fraction was further applied to the Sephadex G15 column (1.6 x 60 cm) in 1% acetic acid to purify the modified nucleoside monitored by a reflection index detector. HPLC with an Agilent Zorbax C18 column (25 x 0.9 mm) was used for the RB69 sample. The chromatography was performed in 0.1% TFA (mobile phase A) and 90% MeCN (mobile phase B) gradient from 3% A for 5 min, then to 100% B in 40 min, with a UV detector at 220 nm. The fractions were pooled and lyophilized before NMR spectroscopy.

### NMR spectroscopy

NMR experiments were carried out on a Bruker AVANCE III 600 MHz (1H) spectrometer with 5 mm Z-gradient probe with acetone internal reference (2.225 ppm for 1H and 31.45 ppm for 13C) using standard pulse sequences cosygpprqf (gCOSY), mlevphpr (TOCSY, mixing time 120 ms), roesyphpr (ROESY, mixing time 500 ms), hsqcedetgp (HSQC), hsqcetgpml (HSQC-TOCSY, 80 ms TOCSY delay) and hmbcgplpndqf (HMBC, 100 ms long-range transfer delay). The resolution was kept <3 Hz/pt in F2 in proton-proton correlations and proton-proton correlations and <5 Hz/pt in F2 of H-C correlations. The spectra were processed and analyzed using the Bruker Topspin 4.1.4 program.

### DLP3 and RB69 thymidylate synthase in vivo activity

The plasmids bearing the ThyA genes were used to transform the T7 Express competent cells (NEB) together with the plasmids harboring the NMPK gene from RB69 or DLP3 to examine the in vivo thymidylate synthase activities. For each transformed strain, a single colony transformant was used to inoculate 0.5 mL terrific medium in a well of a 96-well deep well plate with kanamycin and chloramphenicol antibiotics at the concentrations of 50 and 34 µg/mL, respectively. The 96-well plate was left at 37 °C with agitation until an OD600 reading reached 0.6, then was left to cool to RT before IPTG was added at 0.1 µM final concentration. After IPTG induction, the plate was incubated at 30 °C with agitation overnight. The cultures were centrifuged to pellet the cells and were subjected to plasmid DNA isolation using Monarch Plasmid Miniprep Kit (NEB) following the manufacturer’s protocol. The isolated plasmid DNA was hydrolyzed to nucleosides using the Nucleoside Digestion Mix (NEB) performed at 37°C for >2 hrs. The nucleoside solutions were then filtered through a hydrophilic PTFE 0.2 µm centrifugal filter, where the flowthroughs were subjected to HPLC-MS analysis for examining the nucleoside composition of the 5hoC incorporation. Methods described in the LC-MS section were used for the analysis.

### Computational Methods

Complete bacteriophage genomes were downloaded from the National Center for Biotechnology Information (NCBI; https://www.ncbi.nlm.nih.gov/), and environmental sequences were obtained from the Joint Genome Institute’s IMG/VR metavirome database (24). Genomes and metagenomic contigs were re-annotated using the Domainator software suite (25) executing an automated implementation of the hmmer software suite (26) with Pfam v37 profile HMMs for a reference domain database (27).

Subgenomic regions were visualized and extracted using the Geneious software package (28) and synteny maps of selected gene clusters drawn using clinker (29). Multiple sequence alignments were created using MAFFT (30) and protein secondary-structure diagrams were mapped to MSAs using ESPript (31).

Protein structures were predicted with AlphaFold2 (32) from query sequences and automatically generated MSAs (33). Predictive models of substrate bound enzymes were generated using Boltz-1 (34) and rendered in PyMol (35). Structure based similarity searches against the AlphaFold Database (AFDB) and the Protein Databank (PDB) were performed using the DALI web server (36) and FoldSeek (37) with AF2 generated models as queries.

## RESULTS

### DLP3 and RB69 DNA are highly resistant to restriction endonuclease cleavage, attributed to DNA hypermodifications

Previous research into two closely related *Acinetobacter* phages, vB_AbaM_DLP1 (DLP1) and vB_AbaM_DLP2 (DLP2), with which share high nucleotide sequence identity with DLP3, suggesting the phages have modified DNA (22). It was shown that DNA derived from DLP1 and DLP2 is resistant to digestion by 15 of 16 restriction enzymes tested. Only NdeI is able to digest DLP1 and DLP2 genomic DNA under extended incubation conditions, suggesting a large DNA modification that causes steric hindrance to the enzyme. To gain a better understanding of the possible type of DNA modification these phages have, research into other *Straboviridae* phages with unusual DNA modifications was undertaken. This revealed the *E. coli* phage RB69, which possesses a large dC modification originally identified as arabinosylated 5-hydroxymethylcytosine, although a definitive structure and metabolic origins of the arabinosylated deoxycytidine derivative has not been reported (38). Examination of RB69 and DLP3 genomes revealed both phages lack homologs to the T4 alpha- and beta-glucosyltransferases, which add alpha or beta glucose moieties to 5-hydroxymethylcytosine (39). However, it was found that RB69 and DLP3 encode an arabinose 5-phosphate isomerase, sharing 56.7 % amino acid identity. Given the shared encoding of this enzyme and the prior discovery of an arabinose-based DNA modification in RB69, further investigation was warranted to uncover the specifics of the DNA modification in these phages, such as how the modification may protect the DNA from endonucleases.

To study the protection provided by the DNA modification, genomic DNA was isolated from purified virions of DLP3 and RB69 and challenged with nine Type II restriction enzymes previously tested against T4 DNA and other pyrimidine-modified phages (11). As seen in Figure 1, panel A, DLP3 and RB69 gDNA exhibits resistance to four restriction enzymes. The enzymes NdeI, SwaI, AseI, and ApoI display activity against DLP3 and RB69 gDNA; though surprisingly, DLP3 is resistant to MluCI, which showed cleavage against RB69 DNA (Figure 1 A). The in-silico digests of each genome against the restriction enzyme panel are represented in Figure 1 B, which shows both phages possess the target sequences required for activity of all the enzymes used, suggesting the phages have a large DNA modification capable of protecting the DNA from digestion. Taken together, the results from the restriction enzyme digests suggest the presence of a modification capable of inhibiting most of the endonucleases tested; thus, we turned to analytical techniques to examine the DNA more closely for modifications.

**Figure 1:**
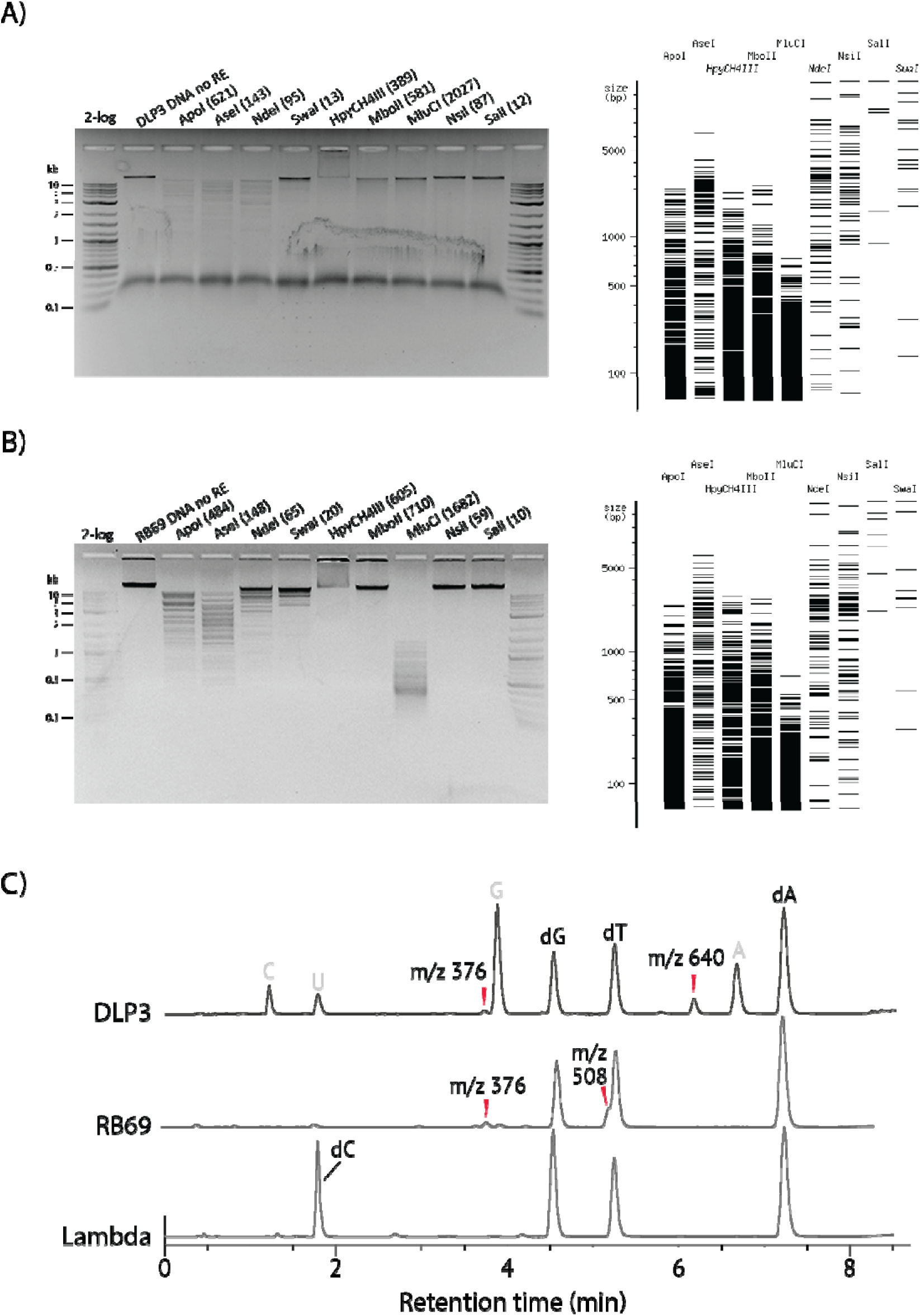
Restriction digests of DLP3 and RB69 virion DNAs and LC-MS analysis of the hydrolyzed DNA for the nucleoside chemical compositions. DLP3 and RB69 virion DNAs were treated with selected restriction endonucleases (RE) and analyzed by agarose gel electrophoresis. The theoretical number of cut sites predicted by REBase on DNA sequences are shown in parentheses next to the given enzyme. Virtual restriction digestion of DNA to show the theoretical digestion patterns in the absence of modifications using NEBcutter tools. A) REs-digested DLP3 DNA analyzed on agarose gel shows on the left and virtual restriction digestion gel on the right. B) RE digested RB69 analyzed on agarose gel on the left and virtual restriction gel on the right. C) LC-MS analysis of the nucleases- and phosphatases-hydrolyzed DLP3 and RB69 DNA. Hydrolyzed phage λ DNA is the reference to show the retention of canonical nucleosides. DLP3 and RB69 show non-canonical peaks corresponding to the hypermodified bases.

### DLP3 and RB69 contain non-canonical bases in their virion DNA

DLP3 and RB69 DNA purified from the virus were enzymatically hydrolyzed and dephosphorylated to nucleosides and then analyzed using liquid chromatography-mass spectrometry (LC-MS). The UV chromatograms are shown in Figure 1C. Despite the minor ribonucleoside carryover in DLP3 DNA, both hydrolyzed phage DNAs have observed canonical nucleosides dG, dT, and dA at 4.5 min, 5.2 min, and 7.2 min, respectively, in which the mass detection confirmed their assignments. However, we could not detect the presence of dC. Instead, non-canonical nucleoside species were detected with m/z 376 and m/z 640 in DLP3 and species with m/z at 376 and m/z 508 in RB69 (Figure 1C and 1-SI). The result suggests that the dC is modified in the phage DNAs. Further analysis of the LC-MS extracted ion chromatography (EIC) (Figure 1-SI) shows a species at m/z 508 which co-migrates with the dT peak in DLP3 nucleoside chromatography. Nonetheless, the observed ion at m/z 640 in DLP3 DNA is absent in RB69. The observed ions at m/z 376 and 508 species in DLP3 and RB69 DNAs have the same retention time, suggesting DLP3 and RB69 likely contain the same dC modifications. Notably, the same mass differences between species at m/z 376 and m/z 508, and species m/z 508 and m/z 640, are concurrently 132, implying the modifications possess a common monomer moiety, likely amended sequentially on top of an ancestor modification intermediate.

### DLP3 and RB69 DNAs contain poly-arabinosylated 5-hydroxycytosines

MS/MS was performed on the non-canonical nucleoside ions to characterize the modifications, and the spectra are presented in Figure 2. The tandem mass spectrum of the precursor ion at m/z 640.22 contains a fragment ion at m/z 524.17, representing the loss of the deoxyribose moiety upon fragmentation (Figure 2A). The further fragmentation produces the fragment ions at m/z 392.13, 260.09, and 128.05, corresponding to the subsequent loss of three pentose residues. Similarly, the precursor ions shared between DLP3 and RB69, at m/z 508.18 and 376.13, contain fragments at m/z 392.13 and 260.09, respectively, denoting the loss of the deoxyribose. Further fragmentation yields the fragment ions at m/z 260.09 and 128.05, corresponding to the succeeding loss of two or one pentose moieties. Our observation is consistent with the previous report by Thomas, J. A. et al., which revealed that the base modifications contain arabinose(s) (38). However, the fragment ion at m/z 128.05 cannot be attributed to cytosine, which should be detected as m/z 112.05. The result thus indicated that cytosine was hydroxylated proximally relative to the glycosyl moiety. It is well known that base modification can occur on the C-5 of cytosine as 5-methylcytosine (5mC) and 5-hydroxymethylcytosine (5hmC) (40, 41). The masses of methylcytosine (42) and hydroxymethylcytosine (43) do not agree with either of these two previously reported modifications because their protonated ions should be detected as m/z 126.07 and 142.06, respectively (44). We propose that the detected ion at m/z 128.05 is attributed to the hydroxylation of cytosine, presumably at the C5 position, i.e.., 5-hydroxycytosine (5hoC).

**Figure 2:**
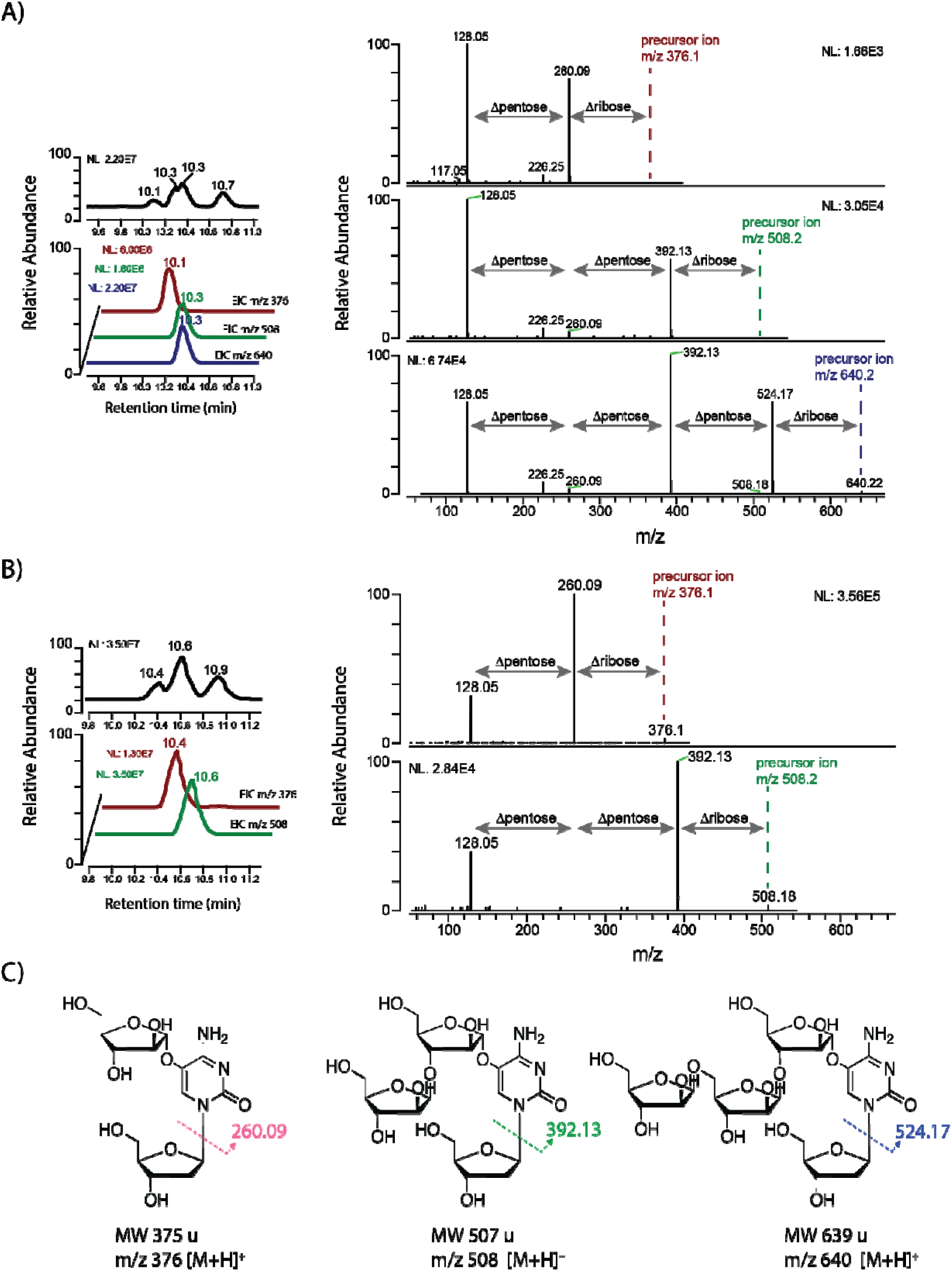
LC-MS/MS analysis of non-canonical nucleosides hydrolyzed from DNAs of *A. baumannii* phage DLP3 and *E. coli* phage RB69. **A)** LC-MS/MS analysis of the hydrolyzed DLP3 DNA. Total ion chromatogram (TIC) and extracted ion chromatograms (EIC) at m/z 640, 508, and 376 are shown on the left and MS/MS spectra of precursor ions of precursor ion at m/z 640.22, 508.18, and 376.13 shown on the right. **B)** LC-MS/MS analysis of the hydrolyzed RB69 DNA. Total ion chromatogram (TIC) and extracted ion chromatograms (EIC) at m/z 508, and 376 are shown on the left and MS/MS spectra of precursor ions at m/z 508.18, and 376.13 shown on the right. C) Chemical structures of the arabinose- modified 5-hydroxy-2′-deoxycitidine non-canonical nucleosides identified in hydrolyzed DLP3 and RB69 DNA were determined using LC-MS/MS and NMR.

We applied NMR to identify the explicit structures of the glycosylated nucleotide hypermodifications. Modified nucleotides were obtained by exhaustive enzymatic digestion and dephosphorylation of the isolated DNA and purified through solid-phase extraction and reverse-phase HPLC. The proposed tri- pentose-modified 5hodC was from DLP3 DNA, and the di-pentose-modified 5hodC was from RB69 DNA. 1H NMR and 13C NMR spectra of the nucleoside isolated were completely assigned using 2D COSY, TOCSY, HSQC, HMBC, and NOESY experiments.

For the tri-pentose 5hodC species, the NMR spectra indicated the spin system of deoxyribose (dR) and three other spin systems of furanose monosaccharides. The three pentoses were identified as _- arabinofuranose (_-Ara*f*) (Figure 2-SI, panel A) and two β-arabinofuranose (β-Ara*f*) (Figure 2-SI, panel B). The configurations of the monosaccharides were determined using NMR chemical shifts in comparison with published data (23)(Table 1-SI). Sequences of the sugars were identified using NOE and HMBC correlations (B1:A3, C1:B5), as shown in Figure 2-SI in Supporting Information. _-Ara*f* A gave no NOE correlations from its H-1, but it gave HMBC correlation to 13C at 128.6 ppm, which correlated to a proton singlet at 7.89 ppm. The signal at 7.89 ppm was the only proton signal of cytosine, identified as H-6 by chemical shift, and consequently, _-Ara*f* A was linked to cytosine C-5. Other cytosine signals were identified using HMBC correlations from cytosine H-6. To verify the identification of arabinose and to determine its absolute configuration, the purified nucleoside and D-arabinose standards were treated with (R)-2-octanol or (SR)-2-octanol and acetyl chloride (10:1, 100 °C, 1 h). The purified sample was dried, acetylated with pyridine-acetic anhydride (1:1, 100 °C, 30 min.) and analyzed by GC-MS (Figure 3-SI). Together, this data identified the structure as β-D-Araf-5-β-D-Araf-3-α-D-Araf-5-hydroxy-2’- deoxycytidine (Figure 2C).

**Figure 3:**
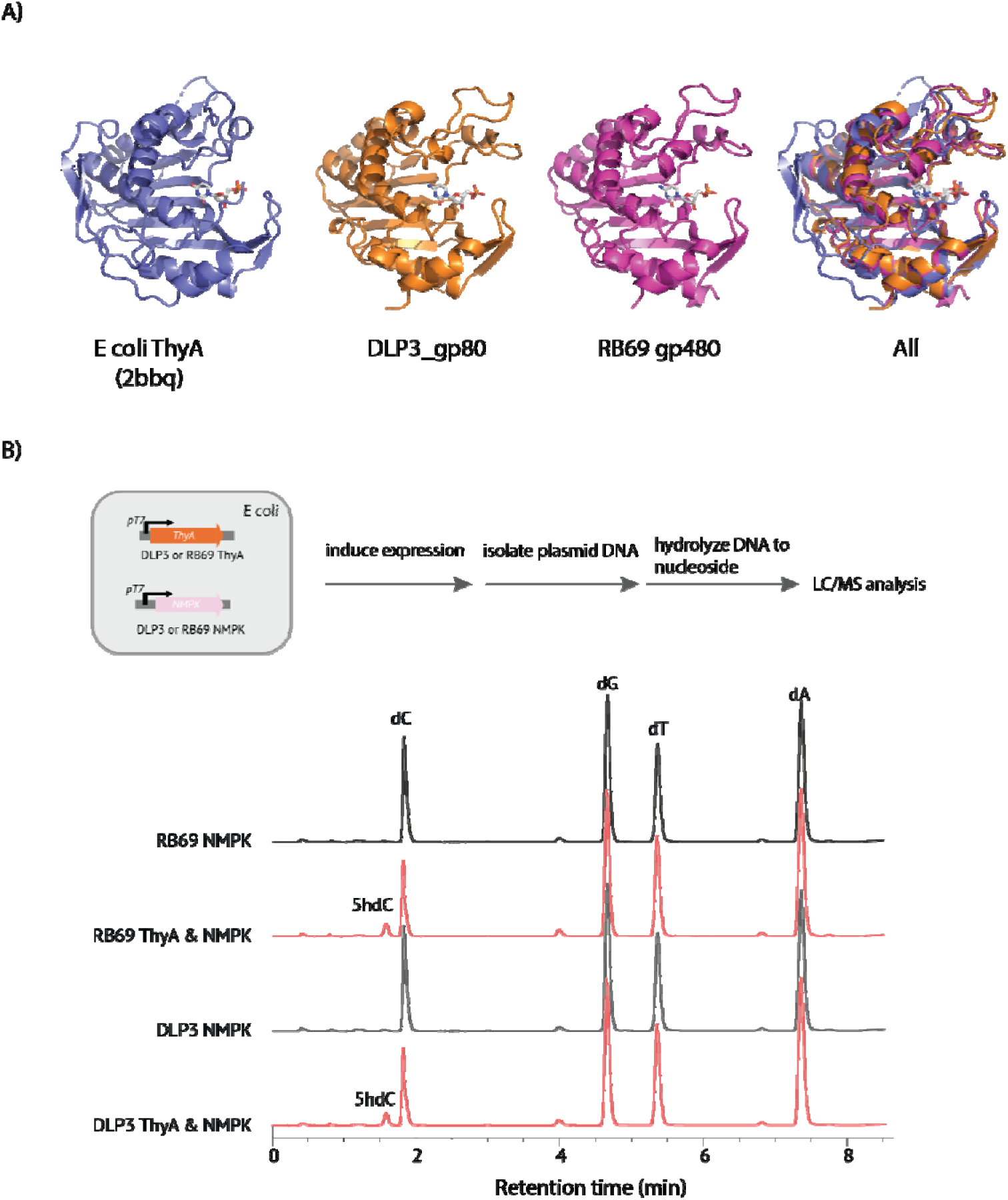
Structure and in vivo activity of thymidylate synthase homologs encoded by phages DLP3 and RB69. **A)** Comparison of homologous structures. From left to right: crystal structure of *E. coli* ThyA bound to dUMP (pdb:2bbq) in blue, predicted dUMP bound structures of DLP3 gp80 (orange) and RB69 gp48 (magenta), and a superposition of all three structures. **B)** Expression of DLP3 gp80 and RB69 gp48, together with their cognate dNMP kinases (DLP3 gp78 and RB69 gp45, respectively) in *E. coli* results in the incorporation of 5hodC into cellular DNA as detected by LC-MS of plasmids recovered from induced cells cultured overnight.

**Table 1.**
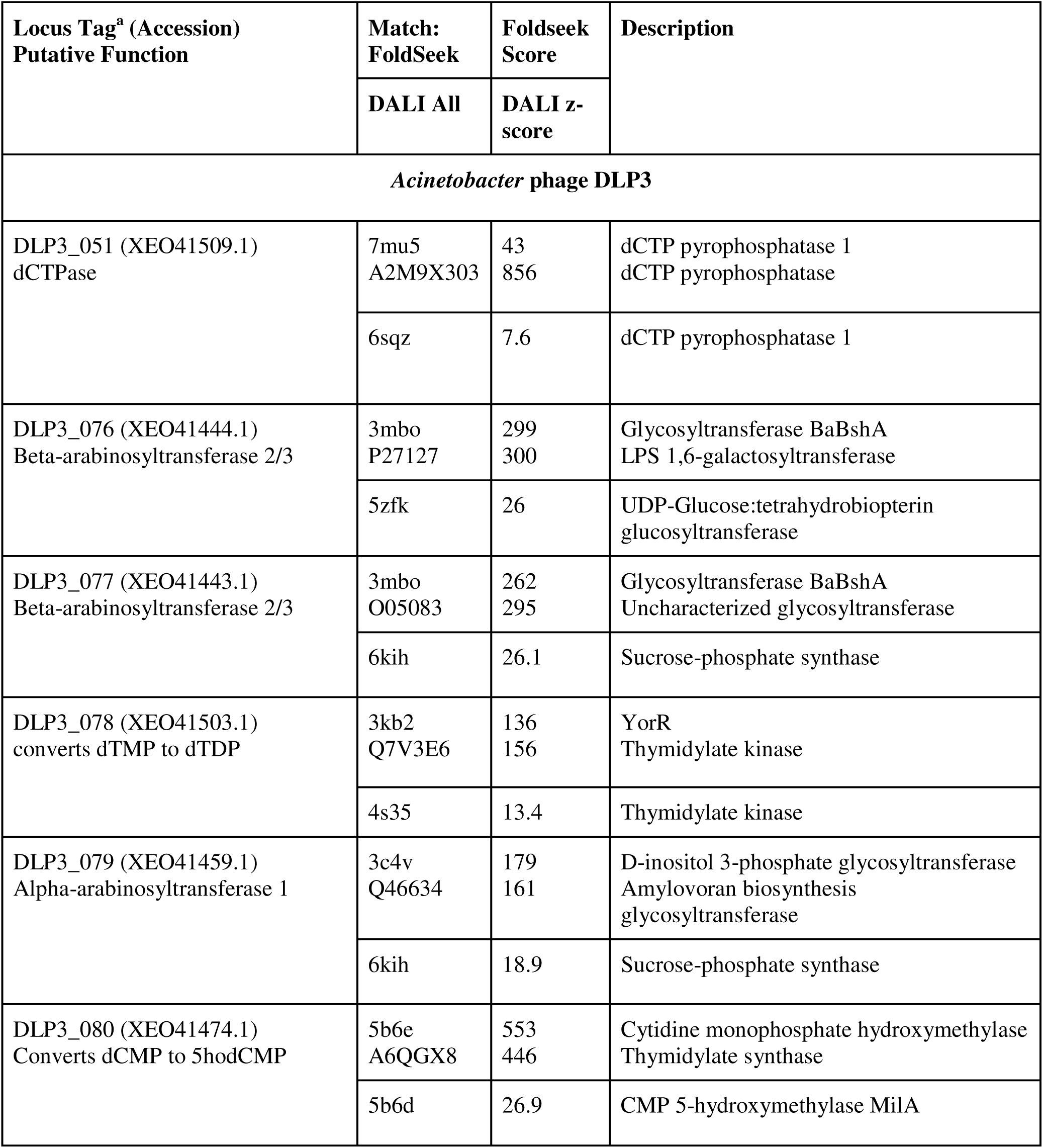
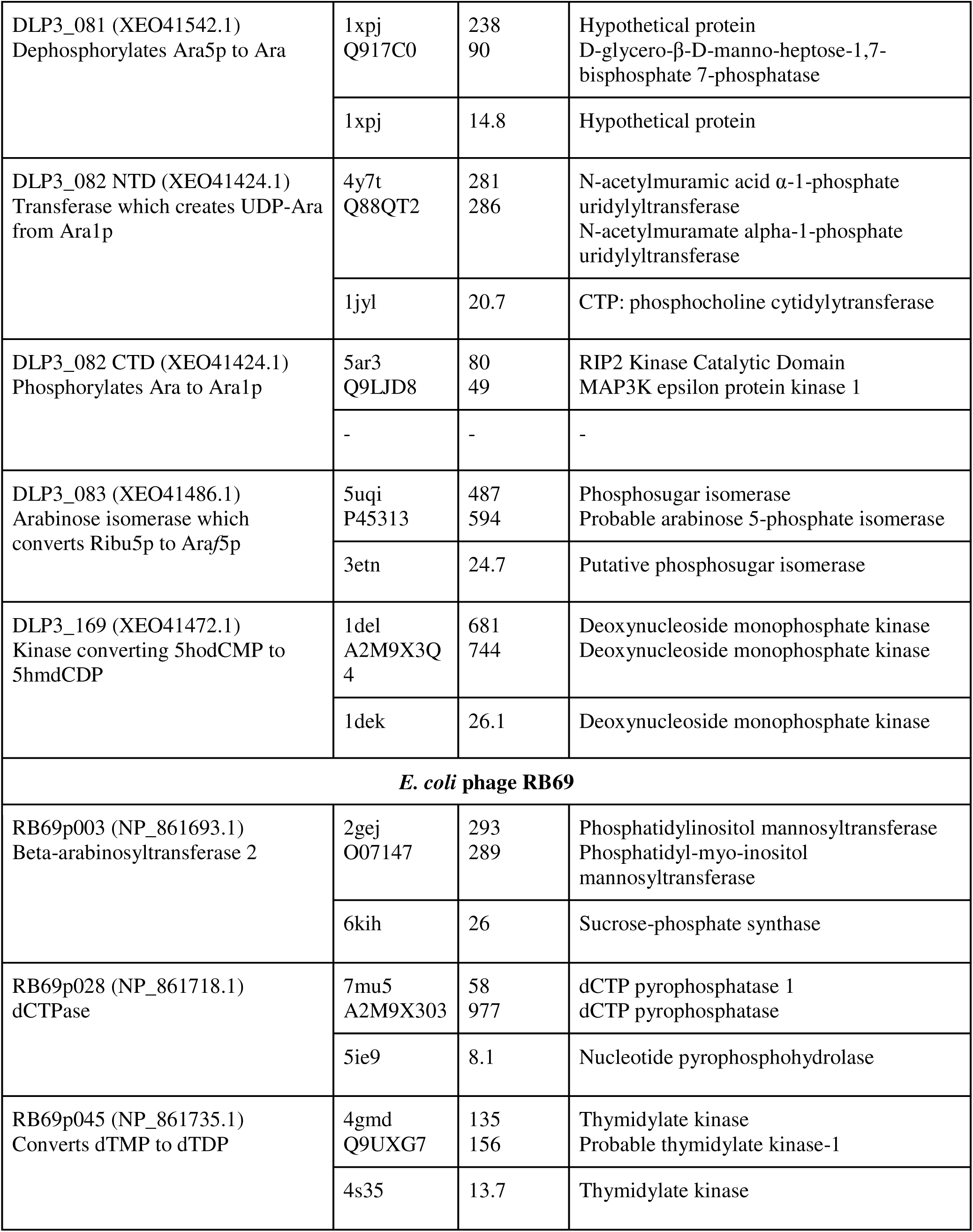
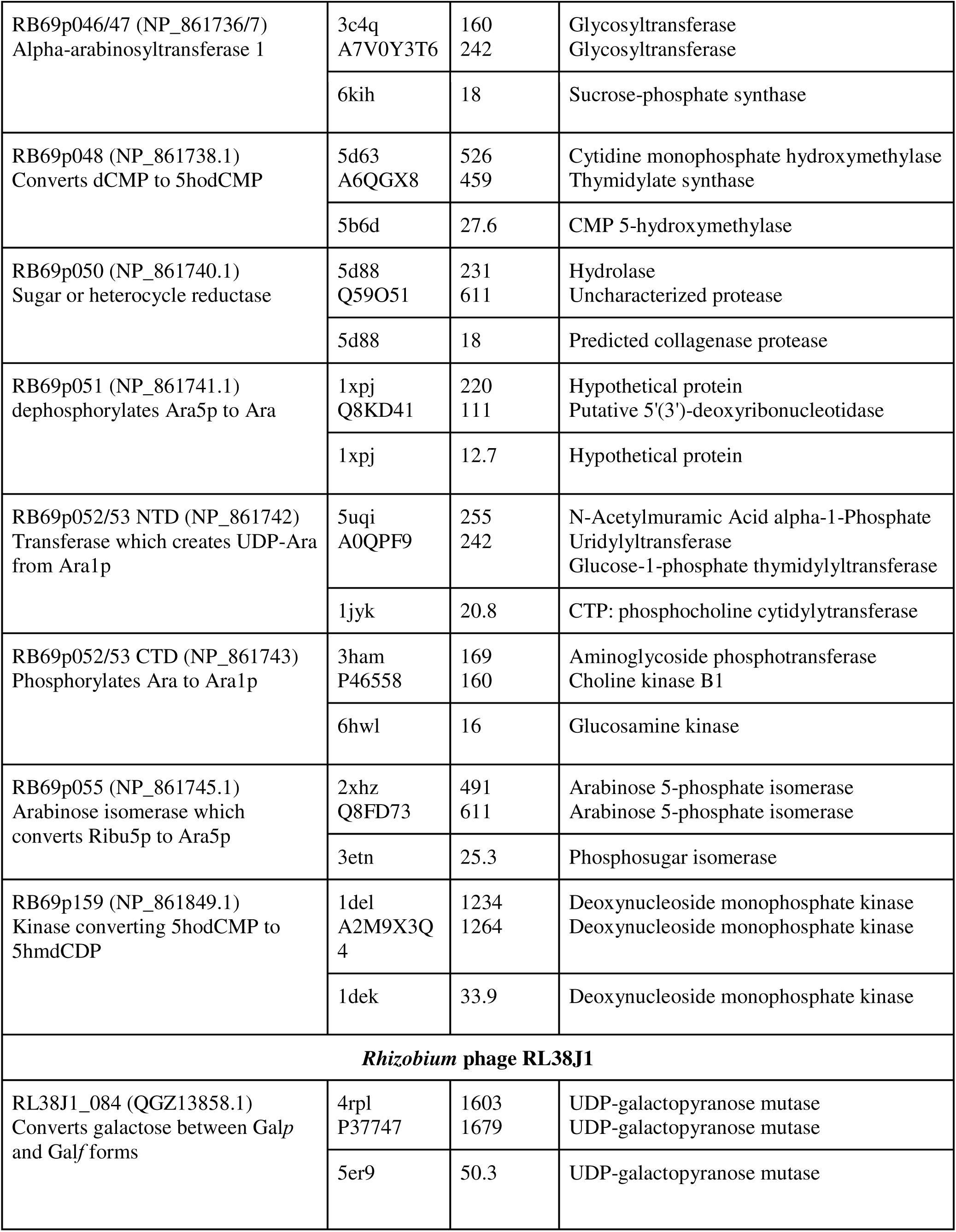
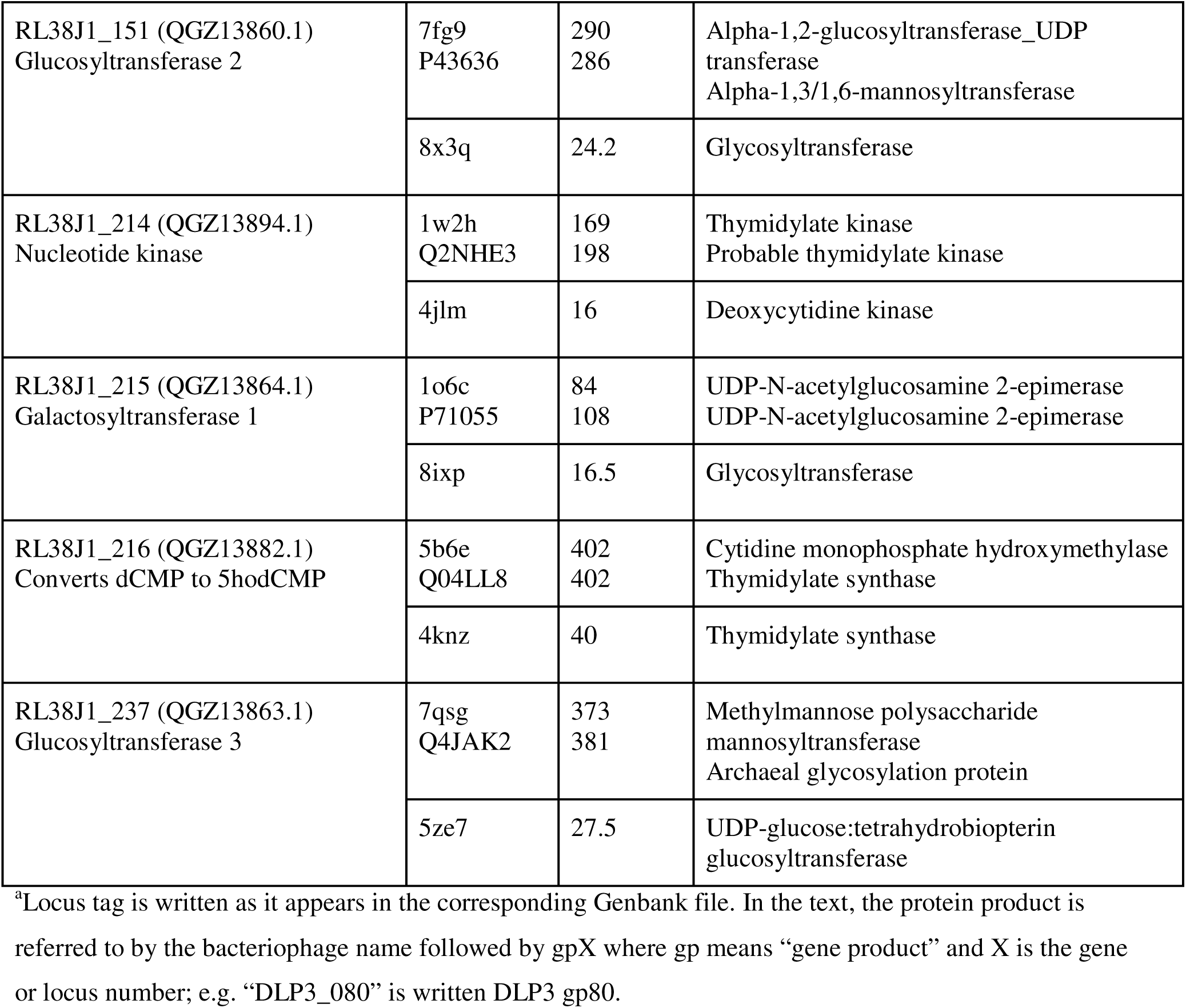
FoldSeek and DALI hits to AlphaFold2 predicted protein structures from DLP3, RB69, and RL38J1.

The di-pentose 5hodC species isolated from RB69 DNA was co-purified with minor dT. Spectra (Figure 2-SI) contained spin systems of deoxyribose (dR) and two other spin systems of furanose monosaccharides, which follow from the low field position of 13C signals >80 ppm for ring carbons.

They were identified as _-Ara*f* (A) and β-Ara*f* (B). Configuration of the monosaccharides was identified using NMR chemical shifts in comparison with the published data (45). Sequence of the sugars was identified using NOE and HMBC correlations (B1:A3). α-Ara*f* gave HMBC correlation A1:N5, which correlated to a proton singlet at 8.05 ppm, the only proton signal of modified cytosine, and consequently, α-Ara*f* was assigned as linked to cytosine C-5. Other cytidine signals were identified using HMBC correlations from cytosine H-6. This data indicates RB69 cytosine is modified to β-D-Ara*f*-3-α-D-Ara*f*-5- hydroxy-2’-deoxycytidine (Figure 2C).

### Biosynthesis of 5hodCMP in bacteriophages DLP3 and RB69

In bacteriophage DLP3, a cluster of genes encoding seven proteins (from gp76 to gp82) is predicted to be involved in the novel DNA modification pathway leading to arabinose-modified cytosines. Similarly, in bacteriophage RB69, the region encoding gp45 to gp55 is demonstrated or predicted to produce enzymes likely leading to the pre-replicative synthesis of 5hodCTP, an arabinosylated nucleotide carrier (NDP- arabinose), and DNA arabinosyltransferases. Preliminary analysis suggests these proteins facilitate the biosynthesis and transfer of an NDP-arabinose precursor, potentially UDP-arabinose, to 5- hydroxymethylcytosine within the phage DNA. Below, we discuss the experimentally verified synthesis of 5hodCMP and the steps leading to its incorporation into DNA, the putative enzymes catalyzing group transfer to DNA of arabinose moieties from a nucleotide sugar carrier to DNA, and the proposed biosynthesis of an NDP-arabinose.

The virion DNA of bacteriophages RB69 and DLP3 contains an arabinosylated 5-hydroxycytosine derivative. Following the established paradigm of T4-like bacteriophages (20), which synthesize 5-hydroxymethyl-2′-deoxycytidine monophosphate (5hmdCMP) from dCMP via a thymidylate synthase homolog, we hypothesized that 5-hydroxy-2′-deoxycytidine (5hodC) biosynthesis in RB69 and DLP3 proceeds through a similar mechanism. Specifically, we posited that a phage-encoded thymidylate synthase homolog converts dCMP to 5-hydroxy-2′-deoxycytidine monophosphate (5hodCMP), which is subsequently incorporated into replicating DNA by the phage DNA polymerase.

Sequence and structural analyses of thymidylate synthases, including *E. coli* thymidylate synthase (ThyA) and bacteriophage T4 5hmdCMP synthase, reveal critical residues within the active site that dictate substrate specificity (Figure 3A). Notably, the residue homologous to N177 in *E. coli* ThyA is crucial for distinguishing between dUMP and dCMP substrates. An aspartate residue at this position (N177D) confers a preference for dCMP (Figure 3B) (46). Furthermore, a bulky hydrophobic residue, such as leucine (L143), at the entrance of the active site cleft in *E. coli* ThyA is thought to exclude water, thereby preventing nucleophilic attack of the exocyclic methylene intermediate and subsequent formation of a hydroxymethyl moiety (5, 47). In T4-like 5hmdCMP synthases, this position is typically occupied by an acidic residue, such as aspartate or glutamate (L143D/E) (Figure 3B). Consistent with Tequatroviruses, the thymidylate synthase homologs encoded by RB69 (gp48) and DLP3 (gp80) possess the N177D and L143E substitutions, suggesting they may utilize dCMP as a substrate and produce a C5 modification other than a methyl group (Figure 3B).

To investigate this hypothesis, genes 48 and 80 from RB69 and DLP3, respectively, were expressed in *E. coli* alongside their corresponding dNMP kinases. Plasmid DNA was isolated from the expression cultures, enzymatically hydrolyzed to free nucleosides, and analyzed by HPLC coupled with mass spectrometry (Figure 3). In addition to the canonical nucleosides, a distinct peak with a molecular weight of 243 Da was observed, consistent with 5hodC. Co-injection with an authentic 5hodC standard confirmed the identity of this peak (Figure 4-SI). These results demonstrate that the thymidylate synthase homologs gp48 of RB69 and gp80 DLP3 can indeed synthesize 5hodCMP from dCMP, thus functioning as 5hodCMP synthases. Comparative genomics across highly syntenic regions from other families among the *Straboviridae* strongly suggests 5hodC biosynthesis extends beyond RB69 and DLP3, including bacteriophages belonging to at least eleven other sub-families of the *Straboviridae* including: Mosigviruses, Bragaviruses, Dhakaviruses, Kagamiyumaviruses, Marfaviruses, Lazarusviruses, Kanagawaviruses, Mosugviruses, Karettaviruses, Tegunaviruses, Tequatroviruses (Figure 4).

**Figure 4.**
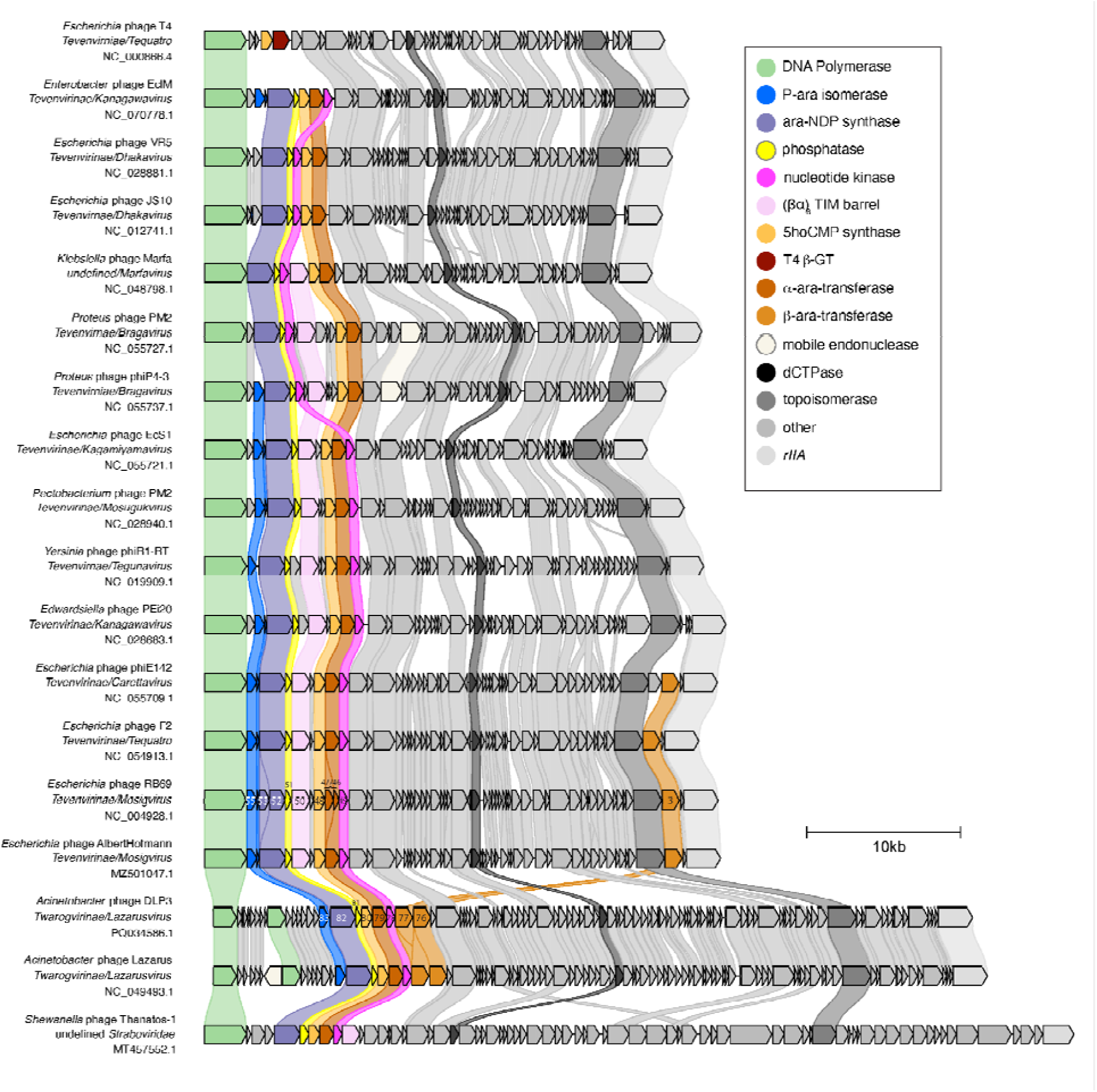
Syntenic organization of cytosine arabinosylation gene clusters across diverse *Straboviridae*. Subgenomic sequences spanning the bacteriophage DNA polymerase to the rIIA gene were extracted from full length genome sequences and annotated using Domainator software suite and mapped using clinker. Genes whose products share greater than 30% amino-acid sequence identity are connected by shaded boxes. Genes are colored according to identified and inferred functions as indicated by the legend in the upper right. Scale bar indicates 10 kb.

Collectively, these bacteriophage groupings infect a diverse range of hosts belonging to the class Enterobacterales, including commensal, pathogenic, and soil dwelling strains.

The hydroxylation reaction carried out by the thymidylate synthase homologs RB69 gp48 and DLP3 gp80 is unexpected chemistry for this class of folate-dependent enzymes. We note that preparations of these enzymes purified by immobilized metal affinity chromatography exhibited a marked yellow color which was lost following additional column purification steps, suggesting a loosely bound cofactor (Figure 5A- SI). UV-Vis wave scans of these preparations yielded absorbance peaks at 370 nm and 455 nm, suggesting a flavin (Figure 5A-SI). HPLC of RB69 gp48 samples treated with protease released a small molecule with identical retention time to flavin mononucleotide (FMN) (Figure 5B-SI). Treatment of the prep with calf intestinal alkaline phosphatase released a molecule with identical retention time to riboflavin, further indicating the identity of the original putative cofactor as FMN or related flavin derivative (Figure 5B-SI). Future work will test what role, if any, a flavin derivative might have in the C5 hydroxylation reaction.

**Figure 5:**
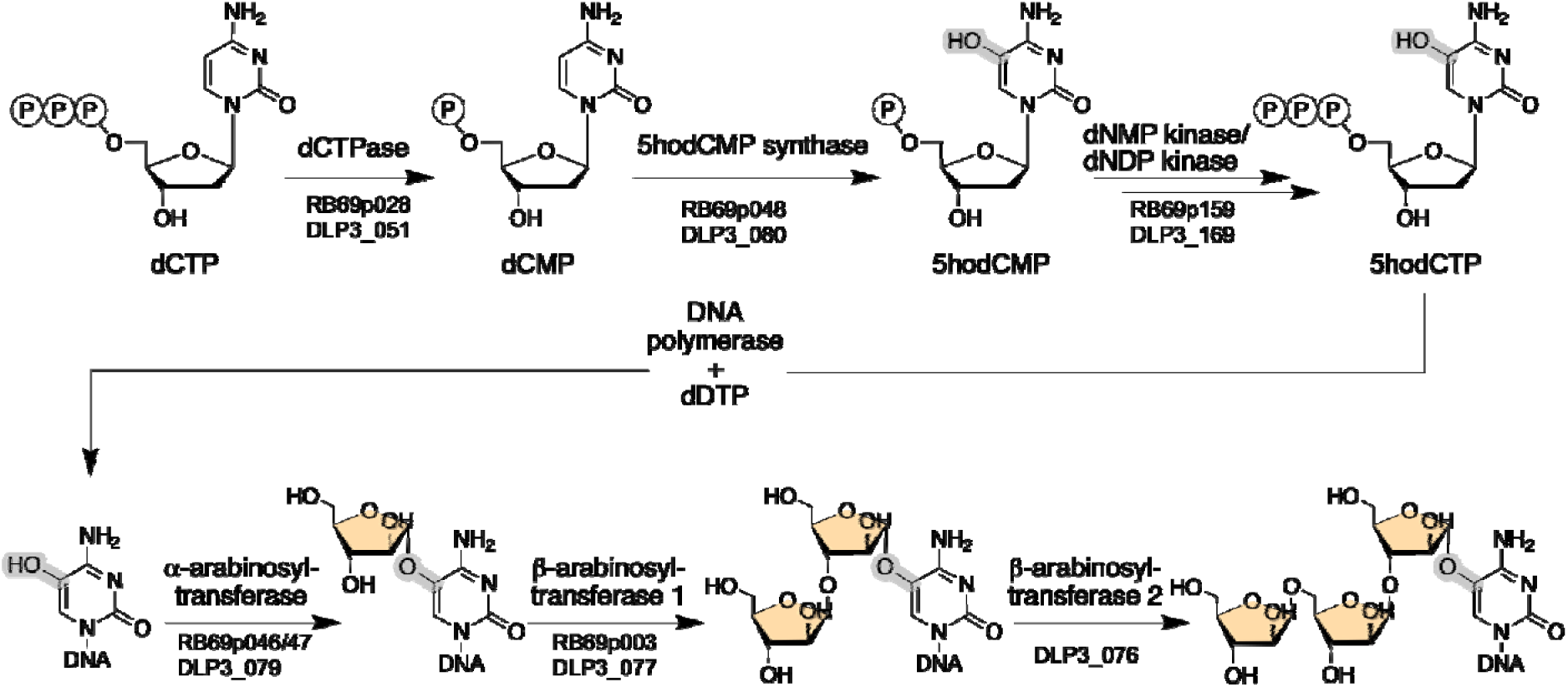
The proposed arabinosylation hypermodification pathway of DLP3 and RB69 DNA. The putative function of each enzyme is listed above the arrow, with the locus tags of each phage encoding the enzyme listed below. The hydroxyl group of 5hodCMP, which is eventually arabinosylated, is highlighted in grey. Both phages encode five enzymes involved in creating and di-arabinosylating 5hodCTP using an alpha- and beta-arabinosyltransferase. Phage DLP3 encodes an additional beta-arabinosyltransferase responsible for the third beta-linked arabinose.

### Proposed Pathway for Arabinose Modification of Cytosine in DLP3 and RB69 virion DNA

To further study the putative enzymes responsible for the transfer of the charged arabinose moiety, we modelled the putative arabinosyltransferases of DLP3 and RB69 using AlphaFold2 and submitted the models to FoldSeek and DALI for structural comparison (Table 1).

Both DLP3 and RB69 encode potential DNA arabinosyltransferases with structural similarities to known glycosyltransferases revealed through FoldSeek and DALI analysis. The putative DNA arabinosyltransferases are encoded by DLP3 gp76, gp77 and gp79 in DLP3, and RB69 gp46/47 and RB69 gp3 (Figure 4). The RB69 genes 46 and 47 are likely a single ORF, but due to a suspected sequencing error, were split into two ORFs; thus, they were concatenated and modelled as one protein for analysis.

Interestingly, the structure prediction models for all the putative glycosyltransferase proteins encoded by DLP3 and RB69 show significant structural similarity to bacterial glycosyltransferases, suggesting a conserved role in glycosylation processes (Table 1).

For DLP3 gp79, the top PDB FoldSeek hit is to a D-inositol 3-phosphate glycosyltransferase (PBD: 3c4v) from *Corynebacterium glutamicum*, which catalyzes the transfer of N-acetylglucosamine from UDP-N-acetylglucosamine to 1-L-myo-inositol-1-phosphate (48). Similarly, DALI results for gp79 and gp77 point to a sucrose-phosphate synthase (6kih) from *Thermosynechococcus elongatus*, which is involved in sucrose-6-phosphate synthesis using UDP-glucose and fructose-6-phosphate (49). The PDB FoldSeek results from the predicted structural models of gp76 and gp77 show significant structural similarity to the glycosyltransferase BaBshA (3mbo) of *Bacillus anthracis*. The BaBshA enzyme produces GlcNAc- malate from UDP-GlcNAc and l-malate (50).

Parallel findings were observed for RB69 proteins. FoldSeek results for RB69 gp3 aligned with phosphatidylinositol mannosyltransferases (PDB: 2gej, AFDB: O07147) from Mycobacterium species, which catalyze the addition of GDP-D-mannose to a lipid carrier (51). The top PDB hit for RB69 gp3 was a N-acetylmuramic acid alpha-1-phosphate uridylyltransferase MurU, which creates UDP-MurNAc from UTP and MurNAc-α1-P in a peptidoglycan recycling pathway (52). DALI results for both RB69 gp3 and RB69 gp46/47 also identified the same sucrose-phosphate synthase enzyme (6kih) as seen with DLP3 gp77 and gp79 (Table 1).

We hypothesize that DLP3 gp79 and RB69 gp46/47 may function as the initial transferase, catalyzing the transfer of arabinose from the NDP-ara to the nucleobase. This hypothesis is supported by the observation that the mono-arabinosylating *Shewanella* bacteriophage Thanatos (21) lacks an ara-GT between the rIIA and topoisomerase genes (Figure 4), suggesting DLP3 gp79 and RB69 gp46/47 may fulfill this role. We propose that RB69 gp3 and DLP3 gp77 may subsequently catalyze the addition of di-arabinosylated species (Figure 5). In the case of DLP3, based on the FoldSeek and DALI structural matches lacking hit similarity to the gp77 and gp79 protein models, we hypothesize the tri-arabinosylated species is added with gp76.

A noteworthy aspect of arabinosylation is the stereochemistry of arabinose attachment. We propose that the configuration around the anomeric carbon (C1) of arabinose may differ depending on whether it is attached to the exocyclic oxygen of 5hodC or to another arabinose residue (likely at C3). Further biochemical and structural studies are required to elucidate these stereochemical details.

### Proposed Pathway for NDP:arabinose biosynthesis in DLP3 and RB69 virion DNA

DLP3 and RB69 both encode three enzymes that could play a role in converting D-ribulose 5-phosphate (Ru5P) to NDP-arabinofuranose (NDP-Ara*f*) for addition to 5hodC: DLP3 gp81-83, and RB69 gp51, 52/53, and 55 (Figure 6-SI, Table 1). We propose that Ru5P is salvaged from the pentose phosphate pathway to act as a substrate for the arabinose 5-phosphate isomerase enzymes encoded by gp83 in DLP3 and gp55 in RB69 to generate D-arabinofuranose 5-phosphate (Ara*f*5P). Both proteins have strong hits to known arabinose 5-phosphate isomerases (53) using FoldSeek and DALI (Table 1), with the DALI search for both proteins hitting to arabinose-5-phosphate isomerase from *Bacteroides fragilis* (3etn).

**Figure 6:**
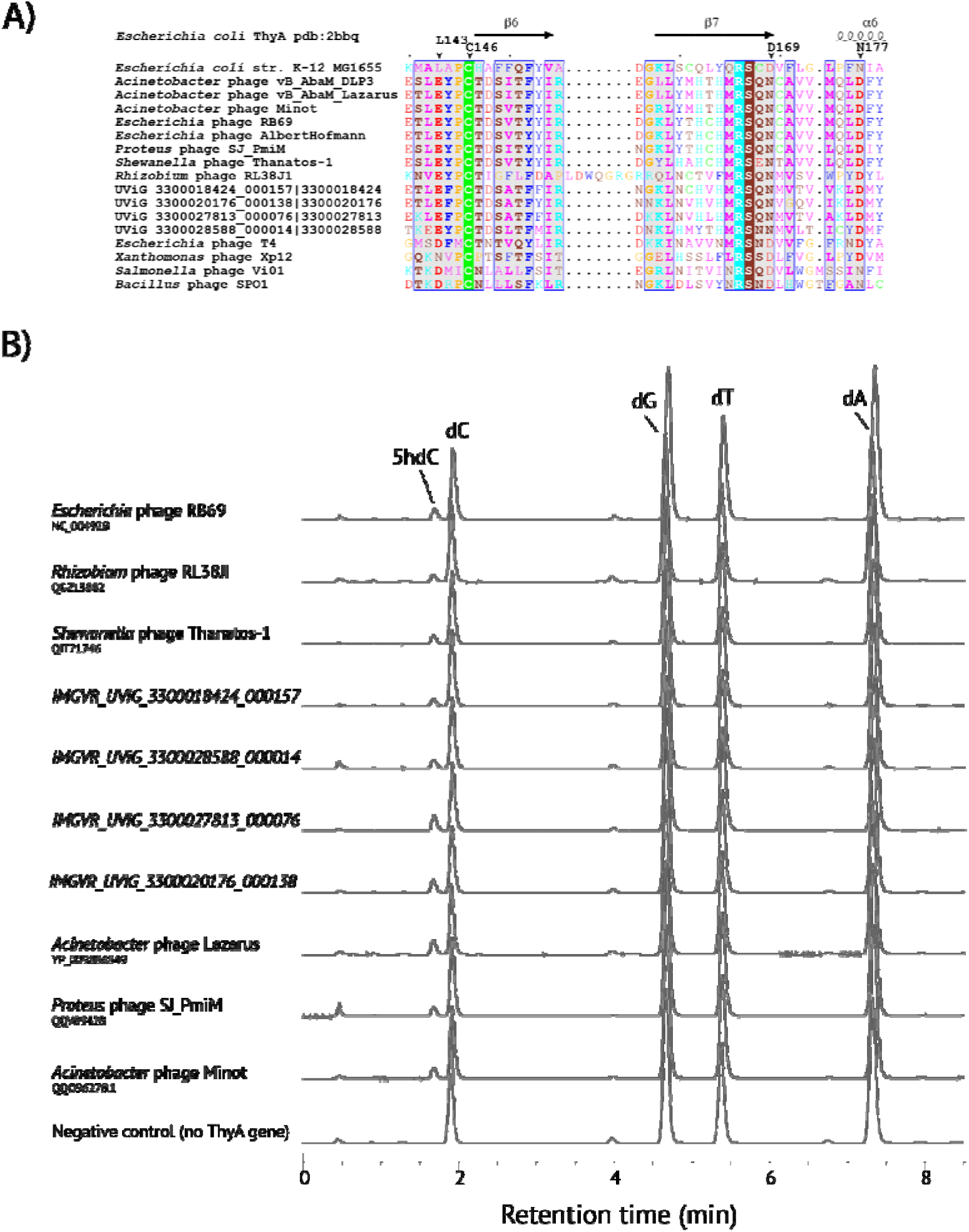
Active site signature and activity of diverse 5hodCMP synthases. **A)** A multiple sequence alignment of residues spanning the active site of *E. coli* ThyA with virally encoded thymidylate synthase homologs of experimentally verified product substrate and product specificities shows replacement of the typically conserved Asp at position 169 with Asn in all homologs tested producing 5hodCMP. Residues are colored by biophysical properties according to the Zappo color scheme. **B)** Experimental validation of 5hodCMP synthesis by naturally occurring D169N variants of virally encoded thymidylate synthases.

The next step in the reaction would be the removal of the 5C phosphate from Ara*f*5P to generate D-arabinofuranose (Figure 6-SI). This step is potentially catalyzed by DLP3 gp81 and RB69 gp51, both of which contain a haloacid dehydrogenase (HAD)-like domain. The HAD-like domain is found among phosphatases, ATPases, beta-phosphoglucomutases, phosphomannomutases, and dehalogenases. The FoldSeek AFDB top hit of DLP3 gp81, and second top hit for RB69 gp51, reveals structural similarity to a D-glycero-beta-D-manno-heptose-1,7-bisphosphate 7-phosphatase (Uniprot: Q917C0) from *Pseudomonas aeruginosa*, which specifically removes the phosphate group at the C7 position of D-glycero-β-D-manno-heptose 1,7-bisphosphate. Thus, these enzymes may dephosphorylate the C5 phosphate of Ara*f*5P to generate D-arabinofuranose for further activation in both phages.

Finally, a nucleoside-diphosphate-sugar pyrophosphorylase could be used to first phosphorylate the D-arabinofuranose to D-arainofuranose-1-phosphate (Ara*f*1P), then convert the Ara*f*1P to NDP- arabinofuranose (Figure 6-SI). A bifunctional enzyme is putatively encoded by DLP3 gp82 and RB69 gp52/53. As with RB69 gp46/47, there is likely a sequencing error that results in a split ORF for RB69; thus, they were concatenated and modelled as one protein for analysis. The FoldSeek and DALI results suggest DLP3 gp82 and RB69 gp52/53 have an N-terminal nucleotidyltransferase domain and a C-terminal kinase domain (Table 1). Both RB69 and DLP3 models have CTD hits to kinases (3ham and 5ar3, respectively), which the phages are potentially using to phosphorylate D-arabinose, creating Ara*f*1P. The Ara*f*1P can then be used by the nucleoside-diphosphate-sugar pyrophosphorylase domain located in the NTD with NTP to generate NDP-arabinofuranose and pyrophosphate. FoldSeek PDB top hits for DLP3 gp82 NTD and RB69 gp52/53 are both to N-acetylmuramic acid α-1-phosphate uridylyltransferases (4y7t and 5uqi respectively). The DALI data supports the FoldSeek findings, with both models having top hits to a CTP:phosphocholine cytidyltransferase from *Streptococcus pneumoniae*. However, based on the structural similarity searches with the predicted models, it is unclear whether UDP-arabinose or CDP-arabinose is the activated sugar molecule used by the glycosyltransferases.

### Diverse viruses encode 5hodCMP synthases and make glycosyl derivatives

Having identified bacteriophage encoded 5hdCMP synthases among diverse members of the *Straboviridae*, we performed multiple sequence alignments of them together with ThyA from *E. coli*. In a ThyA crystal structure (PDB 4isk), atoms from 12 residues can be found within 6 Å of the C5 atom of dUMP: W80, Y94, L143, A142, C146, H147, Q165, S167, C168, D169, G173, and N177 (Figure 6A). Of these, C146 is the catalytic cysteine required for activity (54–56), N177 was previously identified as important in defining substrate specificity (46, 55, 57, 58), and L143 appears to correlate with hydroxymethyl versus methylated product (5, 47). Among the 5hodCMP producing sub-families of the *Straboviridae*, we observed that the amino acid position homologous to Asp169 was consistently occupied by an asparagine; i.e. D169N, relative to *E. coli* ThyA. All other residues were either conserved relative to *E. coli* (e.g. Cys146) or varied within the 5hodCMP producing phages (Figure 6A). Using D169N as a potential diagnostic residue uniquely identifying 5hdCMP synthases, we selected nine diverse thymidylate synthase homologs from metaviromic sequences at the IMG/VR database, distantly related *Straboviridae* (e.g. *Shewanella* phage Thanatos), and the *Rhizobium* phage RL38JI. As shown in Figure 6, panel B, each of these D169N containing thymidylate synthase homologs produced 5hodC in our in vivo activity assay.

The *Rhizobium* phage RL38JI belongs to the *Pootjesviridae*, a family of contractile tail bacteriophages infecting *Rhizobium* and *Agrobacterium* hosts. *Rhizobium* phage RL38JI virion DNA contains a mixture of mono-, di-, and tri- hexosylated cytidines attached to C5 via an ether linkage (59, 60). The base-proximal hexose was identified as galactose followed by one or two glucose units with alpha linkages. Fine grain annotation of the RL38JI genome sequence using HMM profile matching, as well as structural searches using predicted models, finds three CDS features with significant similarity to DNA glycosyltransferases: gp151, gp215, and gp237 (Table 1, Figure 7). Comparative genomic maps generated using clinker on subgenomic regions encompassing these features reveals a high degree of synteny across the *Pootjesviridae*, as seen in Figure 7. The maps suggest a nucleobase hypermodification biosynthetic gene cluster analogous to DLP3 and RB69 with similar components. The proposed hypermodification pathway in RL38JI and related *Pootjesviridae* virions is represented in the Figure 7A-SI, and the biosynthesis of UDP-galactofuranose using a phage-encoded NDP-sugar mutase highlighted in panel B.

**Figure 7.**
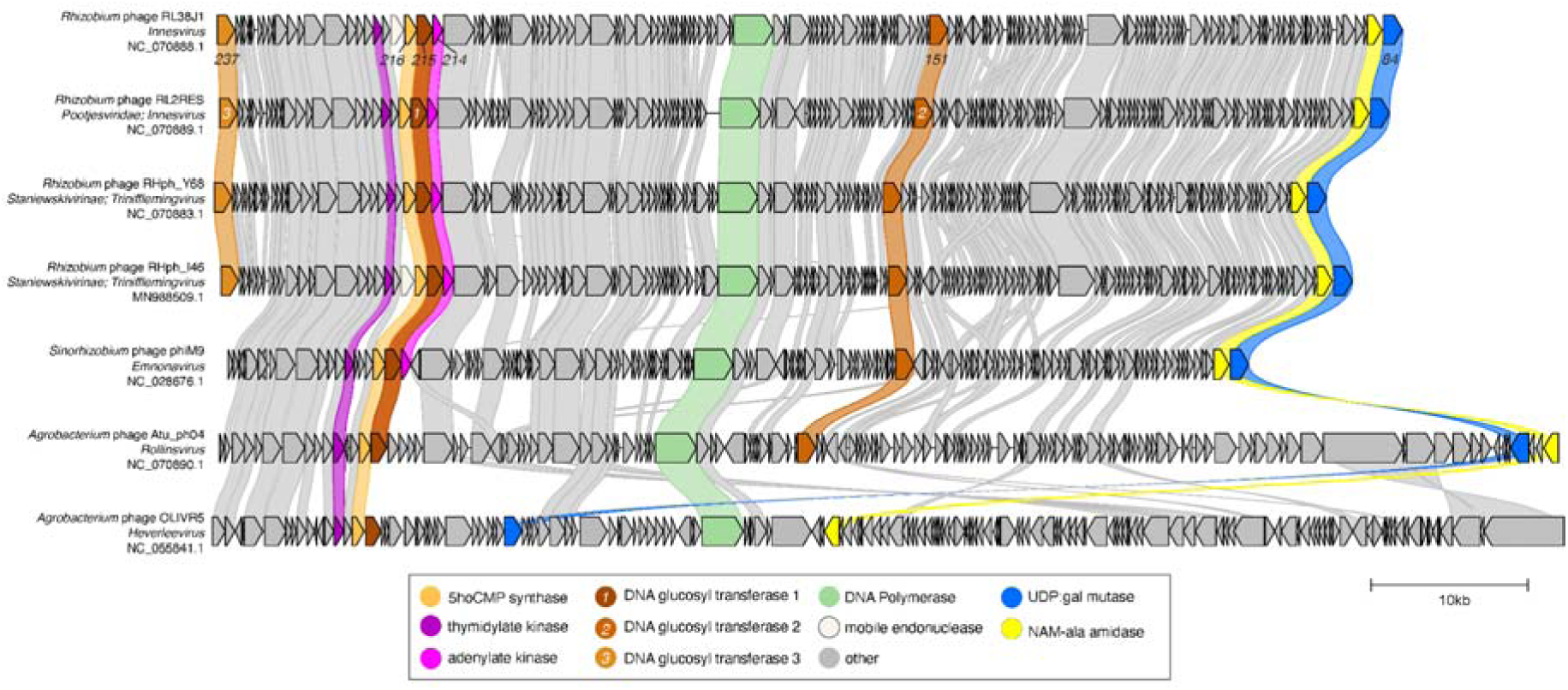
Syntenic organization of DNA glucosylation biosynthetic genes in the *Pootjesviridae*. Subgenomic sequences spanning a syntenic region encompassing DNA glycosyltransferase to a predicted UDP-galactose mutase from representative *Pootjesviridae* were extracted from full length genome sequences and annotated using Domainator software suite and mapped using clinker. Genes whose products share greater than 30% amino-acid sequence identity are connected by shaded boxes. Genes are colored according to identified and inferred functions as indicated by the legend in the upper right. Scale bar indicates 10 kb.

## DISCUSSION

A common theme observed in the biosynthesis of hypermodified bacteriophage DNA is the synthesis of diverse nucleobases by modification of a common set of non-canonical pyrimidines produced at the mononucleotide level by a family of enzymes that includes thymidylate synthase (5, 20, 61). Here we find that multiple bacteriophage families contain cytosines within their DNA to be glycosylated directly at C-5 via an oxygen bridge. We utilized a generalizable in vivo approach to screen the products of thymidylate synthase homologs encoded within these bacteriophage families to uncover an activity not previously seen for these enzymes: cytosine hydroxylation. As with their hydroxymethylated counterparts 5hmdU and 5hmdC, the hydroxyl moiety of 5hodC serves as an attachment point for different chemical groups leading to diverse modifications from a common nucleotide intermediate. The existence of mono-, di-, and tri- glycosylated forms of 5-hydroxycytidine utilizing pentoses or hexoses, suggests other possible combinations of sugars may yet be found to utilize 5hodC as a starting point. The highly conserved amino acid signature of D169N (i.e. the homologous position according to *E. coli* ThyA) can be used to predict thymidylate synthase homologs producing 5hodCMP and will aid in the discovery of other glycosylated forms of cytidine. It should be noted that two other pyrimidine hydroxylation pathways are known to exist, both occurring during the biosynthesis of tRNA hypermodifications (62, 63), but neither of which utilize homologs to the 5hodCMP synthase revealed here. Future work will be aimed at understanding the chemistry of the cytosine hydroxylating branch of the thymidylate synthase superfamily. These discoveries will help illuminate the role of these modifications in the struggle between virus and host as well as inform the selection of bacteriophages for therapeutic applications.

Phages use DNA modifications to protect their DNA from CRISPR-Cas targeting (12, 64, 65) and restriction endonucleases (10, 11) in the bacterial host. The extensive DNA modification present in DLP3 and RB69 DNA was shown to prevent a number of restriction enzymes in the challenge panel from degrading the DNA, despite the presence of their target sites. These findings were similar to data published on T4 gDNA challenged with restriction enzymes, which showed full resistance to MboII, Nsil, and SaII, and partial resistance to SwaI, AseI, HpyCH4III (11). Interestingly, DLP3 DNA showed resistance to the MluCI enzyme, while RB69 DNA was partially sensitive to it. The MluCI enzyme targets AATT sites and is not blocked by methylation. A possible explanation for the differences in MluCI digestion is the tri-arabinosylation of DLP3 5hodC compared to the di-arabinosylation found in RB69. Although the target site is unmodified in both phages, the additional arabinose moiety in DLP3 may create sufficient steric hindrance to shield the target sequences, preventing the activity of the MluCI enzyme against DLP3 DNA.

Unlike Type II restriction enzymes, which can be blocked by DNA modifications, Type IV restriction systems have been discovered and characterized for their ability to target modified DNA, such as GmrSD (14, 66) and McrBC (16). The Type IV restriction enzymes can specifically recognize and cleave DNA modifications such as α-glucose-HMC, β–glucose-HMC, gentiobiosyl-HMC, and other sugar-modified- HMC DNA (16, 66). These types of systems do not target unmodified DNA, which makes them unique in their specificity.

To defend against Type IV restriction systems, phages with modified DNA often encode proteins to block these modification-targeting enzymes. For example, the internal protein I (IPI) is encoded by T4 and is processed into the IPI* form before being loaded into the phage capsid for injection into the host bacterium (67). The IPI* protein has been shown in vitro to interact with the GmrS and GmrD proteins to block their nuclease activity against the modified T4 DNA (68, 69). The RB69 phage encodes an IPI homolog (RB69 gp155), but interestingly, DLP3 does not. Research into the restriction modification systems of Acinetobacter has mainly focused on the impacts of these systems on transformation efficiency (70). The AB5075 host genome in REBASE, a restriction enzyme database hosted by NEB, shows no Type IV restriction systems are encoded, which may possibly explain why the IPI protein is not encoded by DLP3. While the absence of Type IV restriction systems in the *A. baumannii* host may explain the lack of an IPI protein in DLP3, the evolution of the arabinosylation pathway in these phages suggests alternative defence strategies, potentially rooted in the adaptation of existing cellular pathways or horizontal gene transfer.

There are two independent routes that could explain the evolution of the arabinosylation pathway in these phages. First, 5hdCMP synthase might have evolved to bypass anti-phage systems monitoring the presence of 5hmC in the nucleotide pool/DNA, or the destruction of folate species in the cell. Second, arabinose may be a diversification of a base modification to evade systems targeted to glucosylated 5m- cytidines. The broader question is how O-glycosyltransferase enzymes evolved to work on DNA. We speculate that the origins could be from LPS or antibiotic modifying pathways. As outlined in the results section, many of the structural hits for the putative glycosyltransferases are to enzymes encoded by diverse pathogenic bacteria which utilize them to decorate their lipopolysaccharide. While UDP:glucose is a universal metabolite in eubacteria, NDP:arabinose is not known to occur in Enterobacteriaceae.

Arabinosylation is an important modification that can occur in the LPS structures of some Gram-negative bacteria, such as *E. coli, Salmonella typhimurium, Pseudomonas aeruginosa,* and *Burkholderia cenocepacia* (71, 72). This modification primarily involves the addition of 4-amino-4-deoxy-L-arabinose (L-Ara4N) groups to the lipid A portion of LPS. The L-Ara4N modification serves a protective function by reducing the negative charge of LPS, making the bacterial outer membrane more resistant to cationic antimicrobial peptides and certain antibiotics (72, 73). Elements of this pathway may have been co-opted by bacteriophages to arabinosylate their DNA. However, this type of LPS modification has not been identified in *Acinetobacter* species; thus, DLP3 may have acquired the enzymes responsible for its DNA modification through horizontal gene transfer from another DNA-arabinosylating phage. While the evolutionary origins of arabinosylation highlight the intricate adaptations phages have developed to evade bacterial defenses, these modifications also prompt consideration of their implications in phage therapy, particularly regarding how such heavily modified DNA might interact with the mammalian immune system.

One area that is unexplored is the impact heavily modified DNA may have on the mammalian immune system, as phage therapy is being explored as a potential solution to the growing antimicrobial resistance crisis (74). Pathways involved in sensing single-stranded and double-stranded DNA, and induction of cytokines, are commonly identified in response to phage uptake and processing by eukaryotic cells (75–77). The responses appear to be phage specific, as some phages were shown to trigger anti-inflammatory immune responses, while other phages induce pro-inflammatory responses (77). Toll-like receptor 9 (TLR9), which is a pattern-recognition receptor, plays a crucial role in identifying unmethylated CpG motifs that are common in bacterial and viral DNA (78). Stimulation of TLR9 with phage DNA has been shown to lead to the production of inflammatory mediators and induction of interferon gamma (79). This raises questions on how hypermodified DNA affects TLR9 stimulation, and more broadly, other immune pathways. Researching these DNA modifications is crucial to gain a comprehensive understanding of the host immune response to phage therapy, particularly when the treatment involves phage with heavily modified DNA.

## ACKNOWLEDGEMENTS

The authors gratefully acknowledge Prof. Jim Karam (Tulane University) for generously providing a viable specimen of bacteriophage RB69. Critical feedback on the manuscript was provided by Dr. Lana Saleh and Dr. Katherine O’Toole.

## AUTHOR CONTRIBUTIONS

YJ Lee: Conceptualization, Data curation, Formal analysis, Investigation, Methodology, Validation, Visualization, Writing - original draft, Writing – review & editing. Jianjun Li: Data curation, Formal analysis, Investigation, Methodology, Resources, Validation, Visualization, Writing - original draft, Writing – review & editing. Cecilia Sobieski: Investigation, Methodology, Writing – review & editing. Sophie Young: Data curation, Investigation, Writing – review & editing. Jacek Stupak: Data curation, Formal analysis, Investigation, Methodology, Writing – review & editing. Evguenii Vinogradov: Data curation, Formal analysis, Investigation, Methodology, Visualization, Writing - original draft, Writing – review & editing. Hongyan Zhou: Investigation, Methodology, Writing – review & editing. Wangxue Chen: Funding acquisition, Resources, Supervision, Writing – review & editing. Peter Weigele: Conceptualization, Data curation, Formal analysis, Project administration, Resources, Supervision, Validation, Visualization, Writing – original draft, Writing – review & editing. Danielle L. Peters: Conceptualization, Data curation, Formal analysis, Funding acquisition, Investigation, Methodology, Project administration, Resources, Supervision, Validation, Writing – original draft, Writing – review & editing.

## SUPPLEMENTARY DATA

Supplementary Data are available at NAR online.

## DATA AVAILABILITY

Protein and nucleic acid sequences described and analyzed in this work are publicly available through the databases and accession numbers indicated in the text. All other data are available in the main text or in the Supplementary Materials.

