## Supplementary File for "Biosynthesis of glycosylated 5-hydroxycytosine in the DNA of diverse viruses"

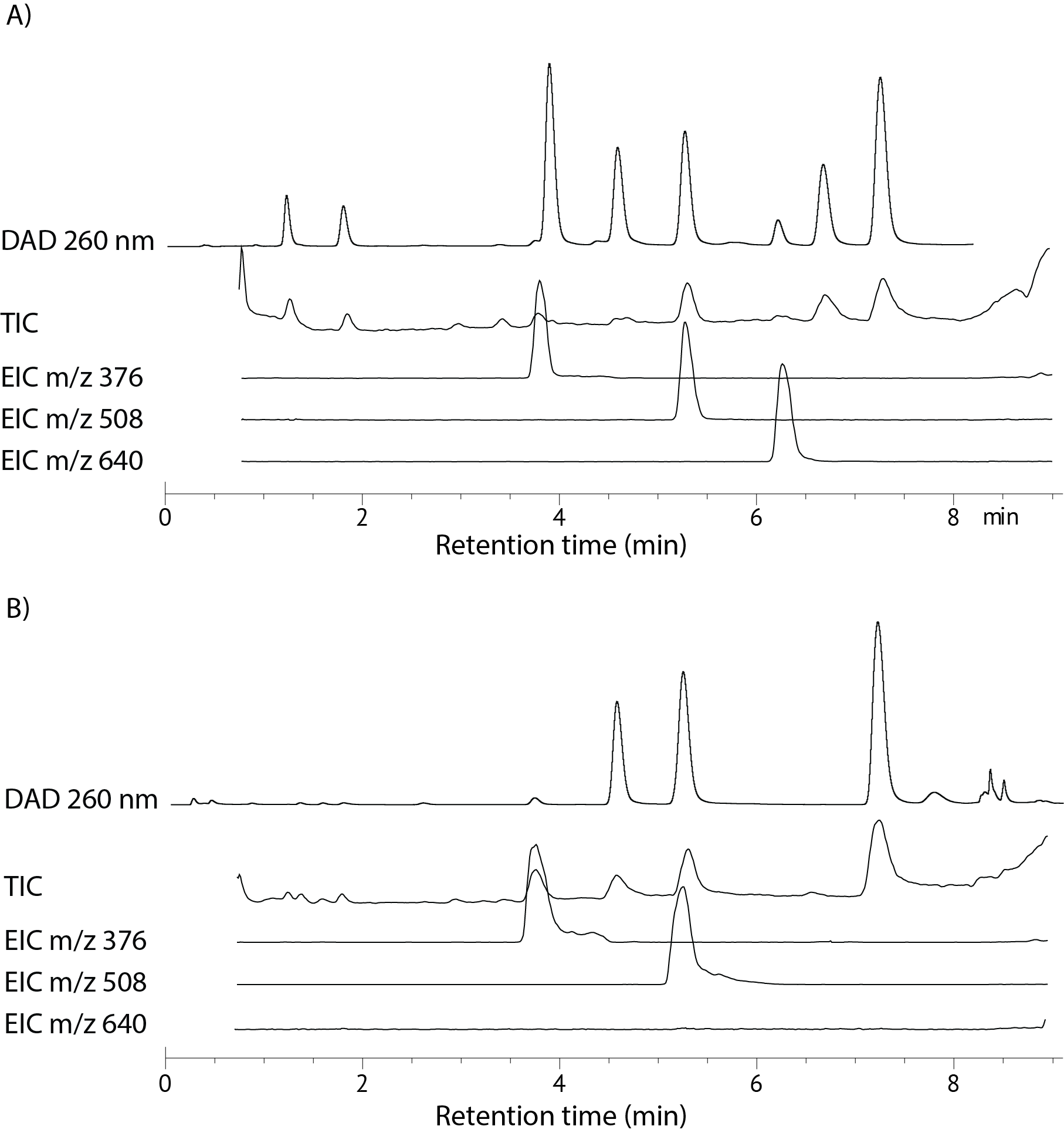

#### **Supplementary Figure 1:** **LC-MS total ion chromatogram (TIC) and extracted ion chromatogram (EIC) of hydrolyzed virion DNA from DLP3 and RB69.** **A)** LC-MS chromatograms of DLP3 nucleoside compositions. The LC-UV chromatogram at 260 nm is aligned with TIC and EIC at *m/z* 376, 508, and 640, representing mono-, di- and tri-arabinosylation of 5hodC, respectively. Three modification products were observed in DLP3 DNA. **B)** LC-MS chromatograms of RB69 nucleoside compositions. The LC-UV chromatogram at 260 nm is aligned with TIC and EIC at *m/z* 376, 508, and 640, representing mono-, di- and tri-arabinosylation of 5hodC, respectively. Mono- and di-arabinosylated products but not tri-arabinosylated products were observed in RB69 DNA.

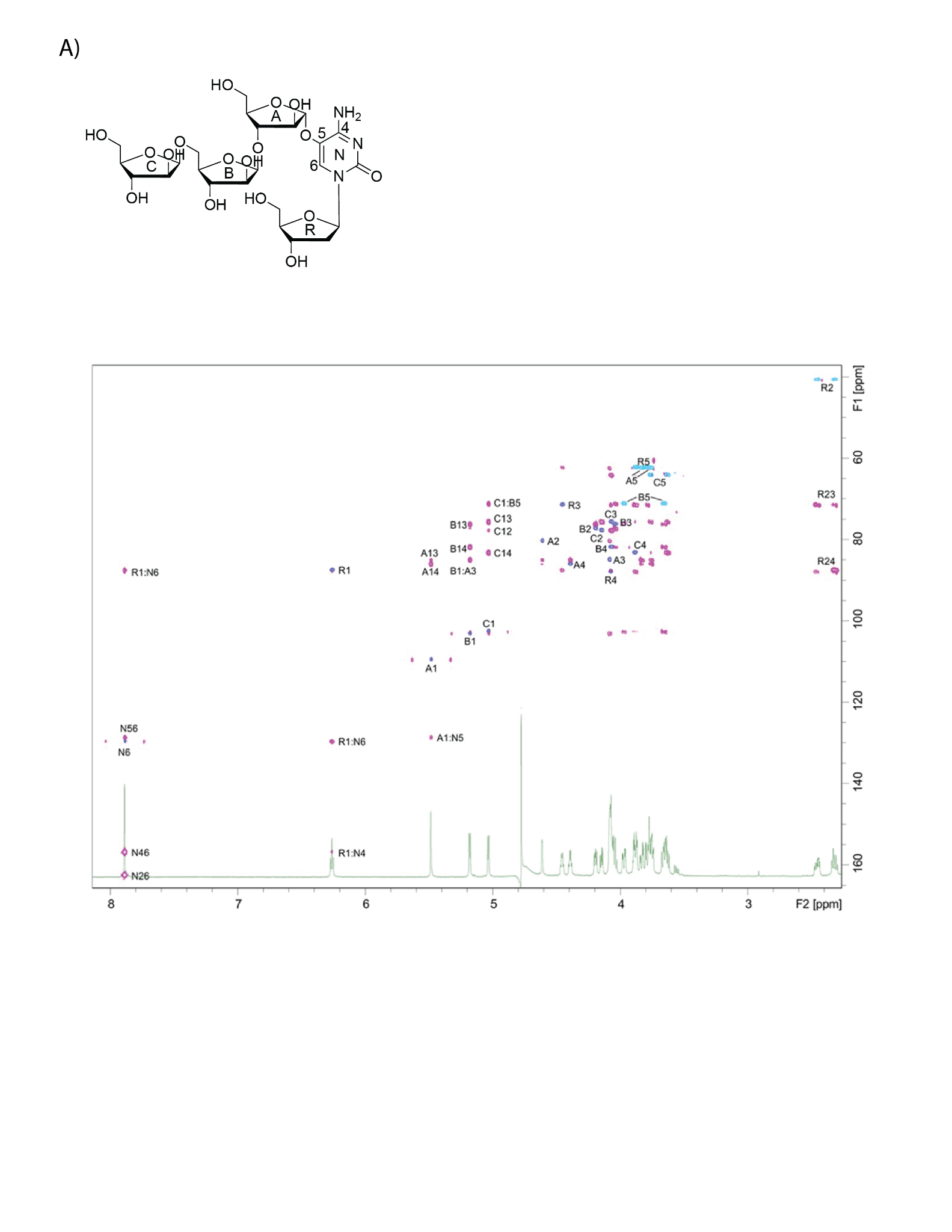

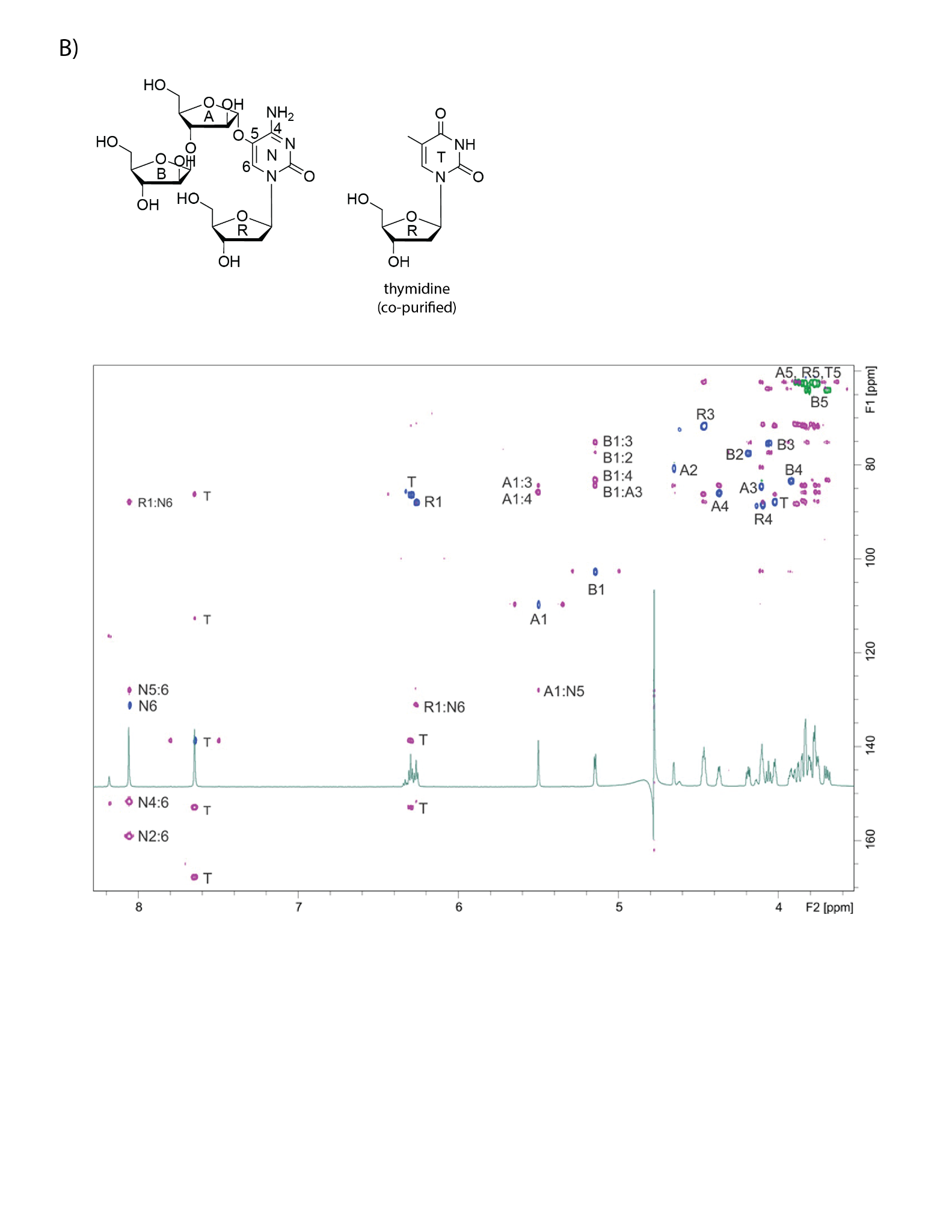

#### **Supplementary Figure 2:** NMR spectra of arabinose-modified 5-hydroxy-2′-deoxycytidine isolated from purified DLP3 and RB69 virion DNA. **A)** ^1^H-^13^C HSQC (blue-green) and HMBC (magenta) spectra of the glycol-nucleoside. R - deoxyribose. **B)** ^1^H-^13^C HSQC (blue-green) and HMBC (magenta) spectra of the OS1 mixture with thymidine. Cytosine label N. Thymidine T.

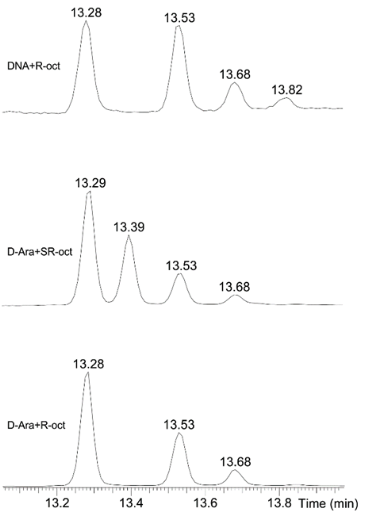

#### **Supplementary Figure 3:** GC-MS analysis of the acetylated derivatives obtained by heating of glycosyl-nucleoside or D-arabinose with (R)-2-octanol or (SR)-2-octanol and acetyl chloride (10:1, 100 °C, 1 h).

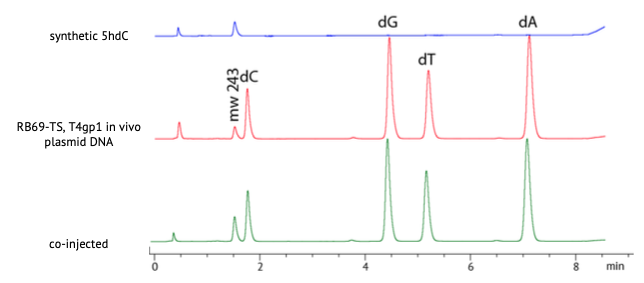

#### **Supplementary Figure 4: Identity of MW 243 nucleoside confirmed by synthetic standard.** Enzymatic hydrolysate of DNA recovered from *E. coli* after overnight expression of RB69 gp48 together with T4 gp1 was resolved by HPLC and mass obtained with MS. HPLC of synthetic 5hodC showed the same mass and retention time as seen in the top trace (blue). Co-injection of the standard with the biological sample resulted in precisely overlapping peaks (bottom trace in green).

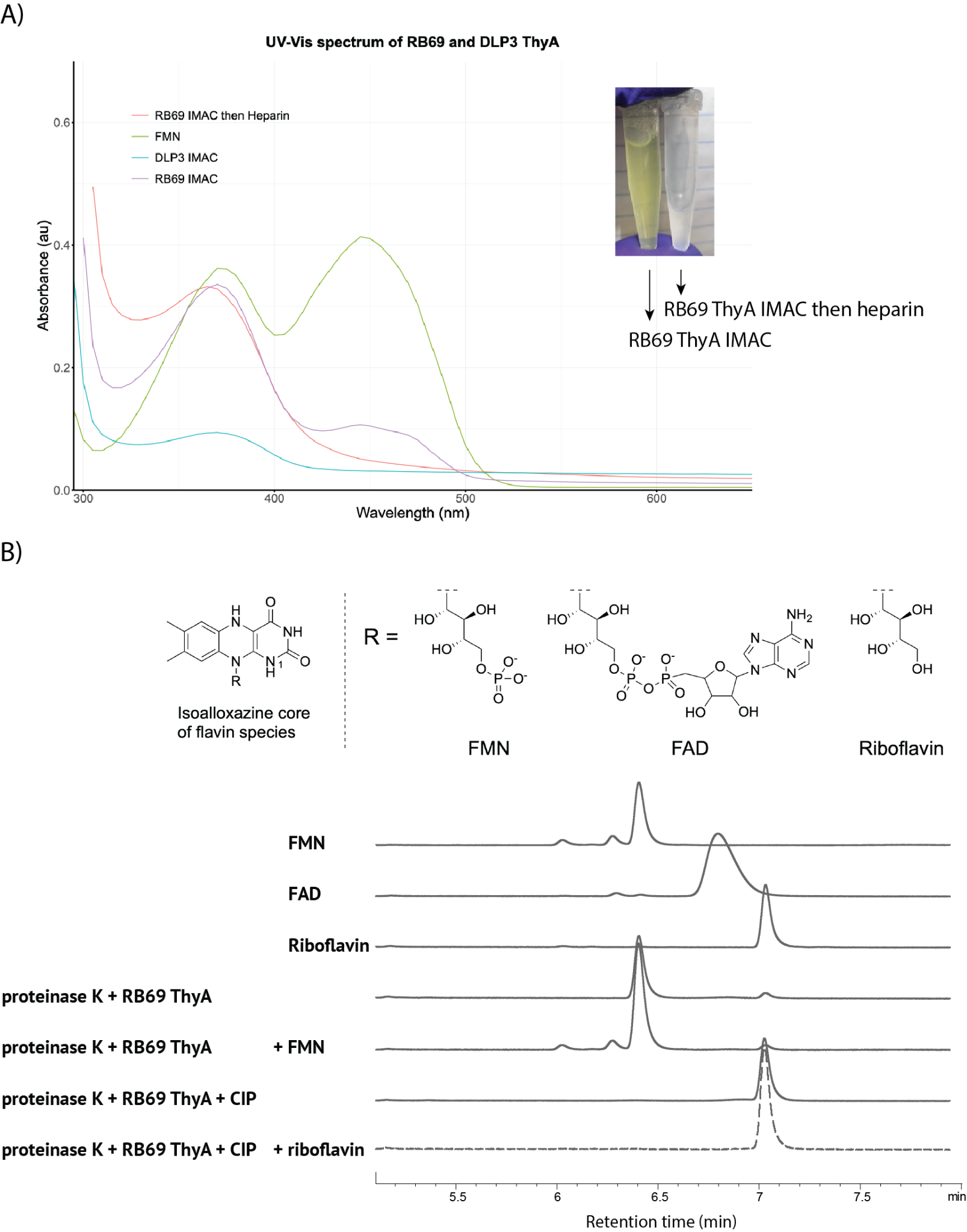

#### **Supplementary Figure 5: UV-Vis spectra of recombinant ThyA proteins from RB69 and DLP3 and the HPLC-UV analysis of the RB69 ThyA protein-bound flavin species.** **A)** UV-Vis spectroscopy scan of the recombinantly expressed and purified ThyAs from RB69 (gp48) and DLP3 (gp80). The ThyA proteins were designed to contain the N-terminal 6xHis tag and were recombinantly expressed and purified through immobilized metal affinity chromatography (IMAC) using Ni-NTA resin. The IMAC-purified protein has a yellow hue visible to the naked eye. The UV-Vis spectrometry was performed on a Costar UV-transparent microplate in a plate reader and the UV-Vis spectrum of the IMAC-purified RB69 ThyA shows two absorbance peaks at ~370 nm and ~455 nm. The protein was further subjected to heparin ion exchange chromatography. However, after the heparin chromatography, the protein lost its chromophore and appeared colorless. The UV-Vis spectrum of the heparin-purified RB69 gp48 has absorbance peaks at ~370 nm retained, but ~455 nm was greatly reduced. **B)** To analyze the protein-bound chromophore, the IMAC-purified yellow-colored RB69 ThyA was treated with thermolabile proteinase K to proteolytically hydrolyze the protein and release the chromophore, which was filtered and analyzed by HPLC-UV monitored at 445 nm. The resulting chromophore appears to be flavin mono-nucleotide (FMN), compared to the FMN standard retention property under the same buffer and chromatography condition. Further treating the released chromophore with calf intestinal alkaline phosphatase (CIP) converts it to a product that has the same LC retention property as riboflavin, which further supports the protein-bound chromophore identity as the FMN.

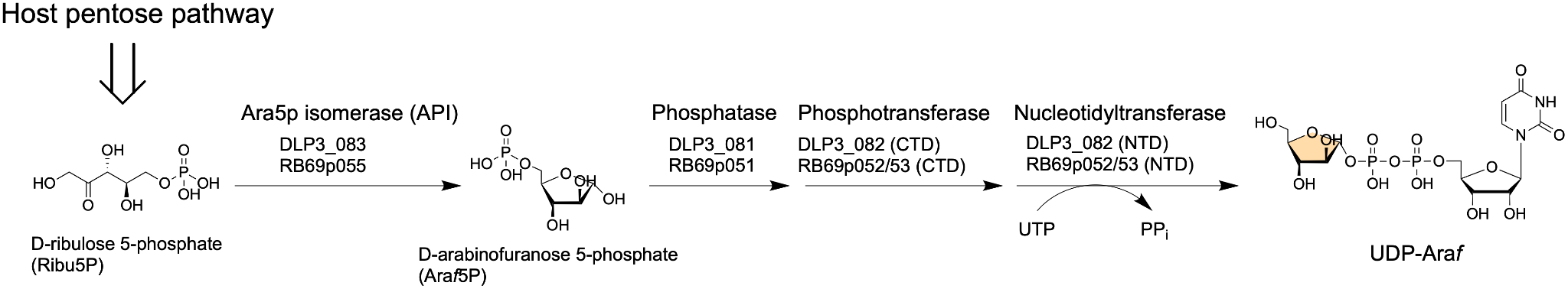

#### **Supplementary Figure 6: Proposed UDP-arabinose biosynthesis pathway in DLP3 and RB69.** The proposed function of each enzyme is listed above the arrow, and the locus tags of each phage encoding the enzyme listed below. Both phages encode three enzymes capable of salvaging D-ribulose-5-phosphate to create the activated UDP-Ara*f* which is used to modify 5hodCTP.

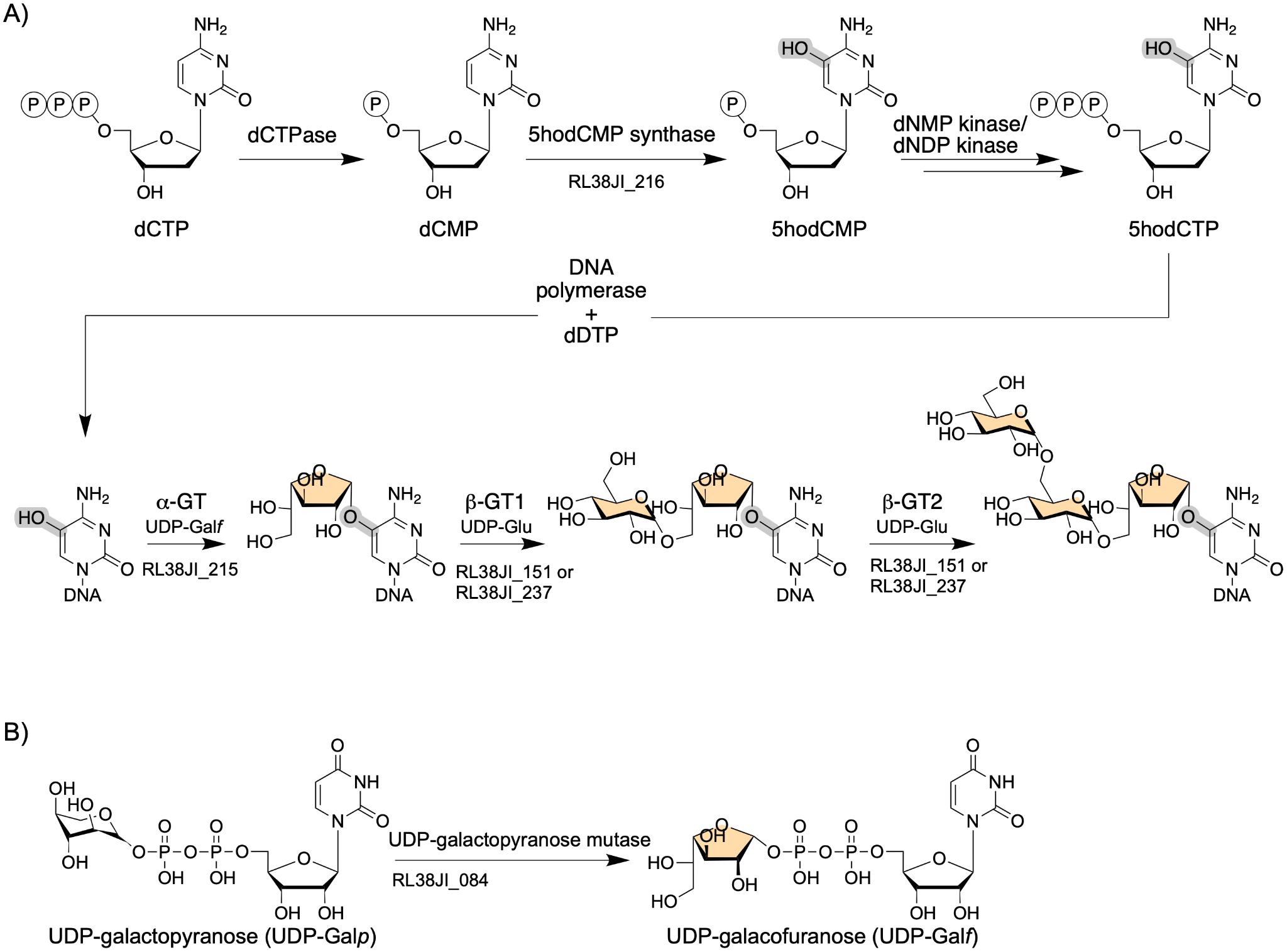

#### **Supplementary Figure 7: Proposed hypermodification pathways of *Rhizobium* phage RL38JI and related *Pootjesviridae*** **A)** Proposed pathway of sugar modification (*i.e.*, (D-Gal)-1-α-D-Glu->6-α-D-Glu) on 5hdC in DNA in the Pootjesviruses. **B)** Proposed biosynthesis of UDP-galactofuranose, the sugar carrier substrate of glycosyltransferase for the initial glycosyltransfer to DNA, isomerized from a putative UDP-galactopyranose by phage-encoded UDP-galactopyranose mutase.

#### **Supplementary Table 1:** NMR data for DLP3 (d, ppm; D_2_O, Bruker 600 MHz 25 °C).

| Residue |  | H/C 1 | H/C 2 | H/C 3 | H/C 4 | H/C 5 | H/C-6 |
| --- | --- | --- | --- | --- | --- | --- | --- |
| α-Ara*f* (A) | H | 5.48 | 4.61 | 4.08 | 4.39 | 3.76; 3.83 |  |
|  | C | 109.5 | 80.2 | 84.9 | 86.0 | 62.3 |  |
| β-Ara*f* (B) | H | 5.17 | 4.19 | 4.04 | 4.07 | 3.66; 3.97 |  |
|  | C | 103.0 | 77.2 | 76.1 | 81.8 | 71.2 |  |
| β-Ara*f* (C) | H | 5.03 | 4.15 | 4.07 | 3.88 | 3.63; 3.76 |  |
|  | C | 102.5 | 77.7 | 75.6 | 83.1 | 64.1 |  |
| Cytosine (N) | H |  |  |  |  |  | 7.87 |
|  | C |  | 162.5 |  | 156.9 | 128.7 | 129.5 |
| 2-deoxyRib (dR) | H | 6.26 | 2.32; 2.45 | 4.45 | 4.07 | 3.78; 3.88 |  |
|  | C | 87.4 | 40.6 | 71.4 | 87.8 | 62.1 |  |

#### **Supplementary Table 2:** NMR data for RB69 (d, ppm; D_2_O, Bruker 600 MHz 25 °C).

| Sugar |  | H/C 1 | H/C 2 | H/C 3 | H/C 4 | H/C 5 | H/C-6 |
| --- | --- | --- | --- | --- | --- | --- | --- |
| α-Ara*f* A | H | 5.50 | 4.65 | 4.11 | 4.37 | 3.76; 3.84 |  |
|  | C | 109.7 | 80.8 | 84.6 | 86.0 | 62.6 |  |
| β-Ara*f* B | H | 5.15 | 4.19 | 4.06 | 3.92 | 3.70; 3.82 |  |
|  | C | 102.7 | 77.5 | 75.4 | 83.3 | 63.9 |  |
| Cytosine N | H |  |  |  |  |  | 8.05 |
|  | C |  | 159.1 |  | 151.8 | 127.9 | 131.1 |
| 2-deoxyRib (dR) | H | 6.26 | 2.36; 2.48 | 4.47 | 4.10 | 3.76; 3.84 |  |
|  | C | 87.9 | 40.9 | 71.8 | 88.5 | 62.6 |  |
